# The Haplotype-resolved Autotetraploid Genome Assembly Provides Insights into the genomic evolution and fruit divergence in Wax apple (*Syzygium samarangense* (BI.) Merr.et Perry)

**DOI:** 10.1101/2023.05.23.542013

**Authors:** Xiuqing Wei, Min Chen, Xijuan Zhang, Yinghao Wang, Liang Li, Ling Xu, Huanhuan Wang, Mengwei Jiang, Caihui Wang, Lihui Zeng, Jiahui Xu

**Affiliations:** Fruit Research Institute, Fujian Academy of Agricultural Sciences, Fuzhou 350013, Fujian, China; Fujian Agriculture and Forestry University, Fuzhou 350002, Fujian, China; Shenzhen Branch, Guangdong Laboratory for Lingnan Modern Agriculture, Genome Analysis Laboratory of the Ministry of Agriculture, Agricultural GenomicsInstitute at Shenzhen, Chinese Academy of Agricultural Sciences, Shenzhen 518120, China

**Keywords:** wax apple (*Syzygium samarangense*), haplotype-resolved autotetraploid genome assembly, transcriptome, fruit size, sugar content, male sterility

## Abstract

The wax apple (*Syzygium samarangense*) is an economically important fruit crop with great potential value to human health because it has rich antioxidant substances. Here, we presented one haplotype-resolved autotetraploid genome assembly of the wax apple with size of 1.59 Gb. Comparative genomic analysis revealed three rounds of whole-genome duplication (WGD) events, including two independent WGDs after WGT-γ. Resequencing analysis of 35 accessions partitioned these individuals into two distinct groups, including 28 landraces and seven cultivated species, and several selectively swept genes possibly contributed to fruit growth, including *KRP1*-*like, IAA17-like, GME-like*, and *FLACCA-like* genes. Transcriptome analysis in three different varieties during flower and fruit development identified key genes related to fruit size, sugar content, and male sterility. We found *AP2* also affects the fruit size by regulating the sepal development in wax apples. The expression of sugar transport-related genes (*SWEET*s and *SUT*s) was high in ‘ZY’, likely contributing to a high level of sugar content. Male sterility in ‘Tub’ was associated with tapetal abnormalities due to the decreased expression of *DYT1, TDF1*, and *AMS*, which affects the early tapetum development. The chromosome-scale genome and large-scale transcriptome data presented in this study offer new valuable resources for biological research on *S. samarangense*, and sheds new light on fruit size control, sugar metabolism, and male sterility regulatory metabolism in wax apple.

## 1. Introduction

Wax apple (*Syzygium samarangense* Bl. Merr. et Perry) also termed Java apple and wax jambu, is a non-climacteric tropical fruit tree from the *Myrtaceae* family and is native to the Malay Archipelago^1^. The *Myrtaceae* family is made up of about 80 genera and 3,000 or more species^2^. According to a few studies of *Myrtaceae* genomes^3,4^, the phylogenetic position remained uncertain. The *Myrtaceae* family have traditionally been divided into two main groups: fleshy fruited and dry fruited^2^. As one of the largest genera of fleshy fruited in *Myrtaceae*, the *Syzygium* species exhibit complex genetic diversity^5^. The *Syzygium* species include *S. aqueum* (water apple, 2*n* = 44), *S. cumini* (Java plum, 2*n* = 66), and *S. samarangense* (wax apple, 2*n* = 33, 42, 44, 66 and 88)^2^. The phylogenetic topologies information based on chloroplast genomes are inconsistent with geographical and morphological classification to some degree^6^. And few *Syzygium* species genomes are available to provide a certain genetic relationship. Accordingly, there is necessary to study the genome information of wax apple to construct a more reliable *Syzygium* species phylogenetic tree. The acquisition of long contigs from autopolyploid or highly heterozygous plants is the major obstacle to obtain accurate genome information, which therefore remains a huge challenge^7,8^.

Wax apple fruit is usually eaten fresh, which is bell-shaped and narrow at the base with four fleshy calyx lobes at the apex. Because of the strong flowering ability, wax apple can fruit in any given season under proper cultivation measures. The fruit has the characteristics of apple-like crispness, the aroma of roses, low-acid taste and rich in antioxidant compounds that are beneficial to human health, and is therefore has become a popular exotic fruit^9,10^. According to statistics from relevant Chinese authorities, the production of wax apple fruit in Taiwan and Hainan provinces was 89,800 tons in 2019 and brought great benefit to local farmers and the country’s economy (data from: http://www.stats.gov.cn/). In order to meet the needs of consumers and enrich the diet with high-quality wax apples with a composition that guarantees high nutritional value, it is important to maintain a suitable sugar content with good size. For some annual crops, *FW* and *POS* gene were identified to modulate fruit size by regulating cell division or expansion in tomato^11,12^, and *CsFUL1* was identified to modulate cucumber fruit size elongation through auxin transportation^13^. However, the genetic information about fruit size regulation in perennial fruit trees is still unclear. In addition, there is low sugar and sour contents in fruit in the most of wax apple varieties. The regulatory mechanism of sugar and acid metabolism in wax apple is also unknown. Therefore, it is need for the genome assembly and whole-genome re-sequencing to further clarify the regulatory mechanism related to fruit quality in wax apple.

It is well known that seedless is an important target trait in fruit breeding. The consumers prefer the seedless trait of wax apples, which were most selected from bud transformation in wild-type. It is a great challenge for breeders to breeding new seedless wax apple cultivars by cross-breeding, and no new cultivars have bred for more than decades. There is still a lack of research on the genetic regulation mechanism of wax apple. Seedless character caused by male sterility has been developed, such as grape, tomato, and citrus. In plants, the male sterility refers to the inability to produce the dehiscent anthers, viable male gametes, and functional pollen. Previous studies have confirmed that the male sterility had two major categories. The male sterility that resulted from the genes both in mitochondria and nuclear was identified as the cytoplasmic male sterility (CMS); the male sterility that resulted from the nuclear genes alone was known as the genetic male sterility (GMS)^14^. For years, wax apple breeding efforts were hampered due to the complex genetic diversity and the lack of genome information. Therefore, an accurate reference genome of wax apple is essential for understanding the mechanisms regulating fertility and accelerating genomic selection breeding efforts.

In previous work, a superior clones ‘Tub Ting Jiang’ (‘Tub’) has been selected, with large and seedless fruit, sweet (total soluble solids ‘TSS’ content is about 10%) and beautiful color^15^. We also collected two special wax apple varieties, ‘DongKeng’ (‘DK’) and ‘ZiYu’ (‘ZY). ‘DK’ is a rootstock variety with rich seeds in all its fruits. And the fruit of ‘ZY’ is bright red, small but high sweet (TSS is about 14%) with 0 to 2 seeds inside. These varieties will be good materials for studying the genome information of wax apple. Through the study on wax apple genome, we hope to accelerate the breeding process and produce more new varieties which are larger, sweeter and more colorful.

In this study, we aim to sequencing and assembly of ‘Tub’, which is an autotetraploid wax apple variety, to fill wax apple genomic information gaps. This genome was used to conduct a comparative genomic analysis to further insight into the functional and structural features of the *S. samarangense* genome. Furthermore, we identified the key genes associated with the fruit size, sugar content and male sterility, which are important breeding traits of wax apple. This genome will provide a valuable resource for further molecular functional analyses and benefit to accelerate breeding of wax apple.

## 2. Results

### 2.1 Genome assembly and annotation

To investigate the feature of the *S. samarangense* genome, we first performed the genome survey analysis, *K*-mer analysis shows multiple peaks at various sequencing coverages, which was consistent with the distribution characteristics of auto-polyploids (**Supplementary Figure 1**). We further validated that it is an auto-tetraploid genome with 44 chromosomes (2*n* = 4*x* = 44) based on 5S rDNA FISH experiment in the karyotype analysis (**Supplementary Figure 2**). The estimated monoploid genome size of *S. samarangense* was 420 Mb with heterozygosity of 1.16% based on the *K*-mer analysis. This is consistent with the evaluation by flow cytometry (1.62 Gb/2C), which contains four haplotypes. To generate a haplotype-resolved genome assembly, we sequenced a total of 92.0 Gb PacBio subreads (∼220 x of the estimated monoploid genome size), 90.0 Gb Illumina short reads, and 92.40 Gb high-throughput chromatin conformation capture (Hi-C) reads (**Supplementary Table 1**). The initial contigs were assembled using the CANU assembler^16^, resulting in a 1.49-Gb assembly with a contig N50 of 304.5 kb (**Supplementary Table 2**). All contigs were further anchored onto 44 pseudo-chromosomes with 11 homologous groups by subjecting to ALLHiC phasing, finally, a total of 1.59 Gb phased assembly sequences were obtained after gap filling, representing an allele-ware, chromosome-scale genome assembly with completeness of 98.9% evaluated by BUSCO (**Figure 1a, Supplementary Figure 3, Supplementary Table 3**)^17^. In addition, approximately 95.6% of the Illumina clean data can be aligned onto the genome assembly, covering 97.9% of the genomic regions (**Supplementary Table 4**), suggesting the high-quality genome sequences were acquired.

**Figure 1.**
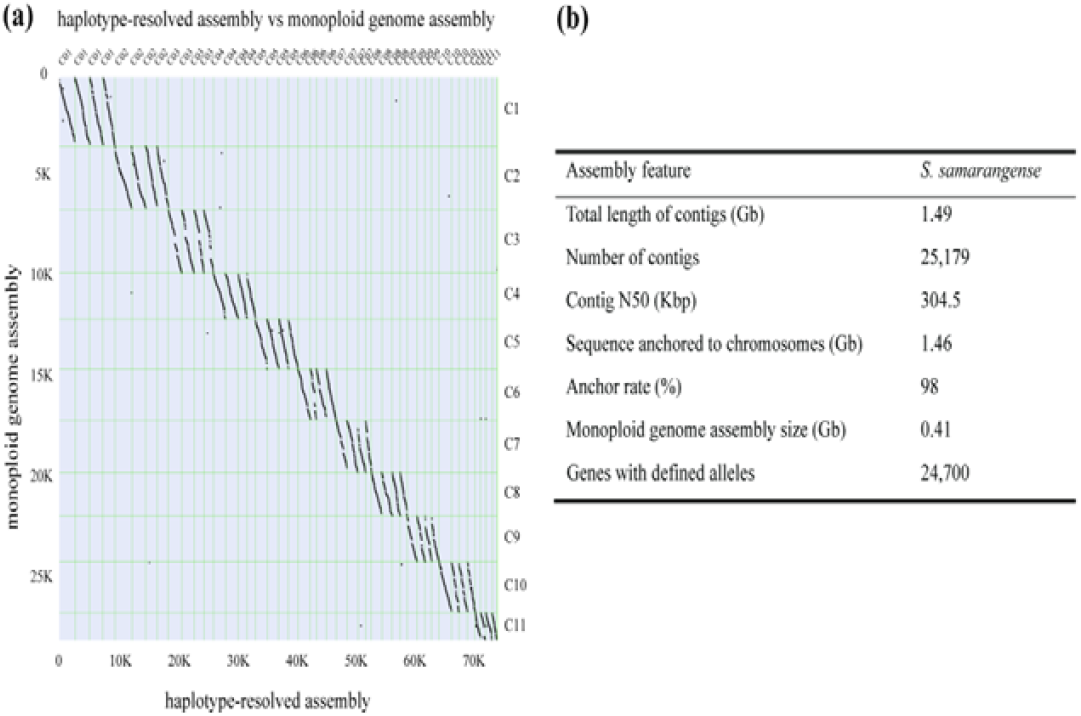
Alignment of *S. samarangense* monoploid genome with *S. samarangense* genome and summary of genome assembly. (a) A set of 4 homologous chromosomes aligned to a single monoploid chromosome. (b) Statics for genome assembly of wax apple.

To gain the high-fidelity gene annotation, we used two rounds of MAKER pipeline to produce a set of 74,888 high-quality protein-coding gene models (**Figure 1b**). BUSCO analysis showed a completeness of 90.7% with 69.3% duplication (**Supplementary Table 5**), indicated that the annotation mixed genes and alleles. We adopted our previously developed pipeline in the sugarcane genome project^18^ to separate genes and alleles, resulting in a total of 24,016 genes with defined alleles. We observed 2,140 (8.9%) genes with four alleles, 7,274 (30.3%) with three, 9,021 (37.6%) with two, and 5,581(23.2%) genes with one. Taken together, our study characterized 52,826 allelic genes, distributed in 24,016 genes with an average of 2.2 alleles per gene. In addition, we annotated 952 tandemly duplicated genes, and 11,161 dispersedly duplicated paralogs (**Supplementary Table 6**).

The wax apple genome contains a moderate level of repetitive sequences (593.25 Mb), accounting for 38.10% of the assembled genome (**Supplementary Table 7**). The long terminal retrotransposons (LTRs) are the predominant transposable elements (TEs) and account for 24.74% of the genome, which consist of 5.76% Ty1/*Copia* and 14.72% Ty3/*Gypsy* (**Supplementary Table 7**). The high proportion of LTRs was likely due to a recent large-scale burst that happened ∼0.1 million years ago (Mya) (**Supplementary Figure 4**).

### 2.2 Evolutionary history and whole-genome duplication

We identified 221 single-copy genes from eight sequenced genomes by OrthoFinder and subsequently employed them to construct a phylogenetic tree. The results clearly presented that *S. samarangense, E. grandis*, and *P. granatum* belong to the same branch of Myrtales. A significant closer genetic relationship was observed between *S. samarangense* and *E. grandis*, which both belong to the *Myrtaceae* family. We further estimated the divergence times and found that Myrtales arose 79.4 million years ago (Mya). Within the *Myrtaceae* family, *S. samarangense* and *E. grandis* diverged from each other at 26 million years ago (Mya). According to a CAFE analysis, we characterized 1,328 gene families expanded and 5,363 under contraction (**Figure 2a**). Gene Ontology (GO) enrichment analysis showed that the 1,328 expanded gene families were majorly enriched in DNA polymerase activity, retrotransposon nucleocapsid, and mitochondrial fission. In contrast, the 5,363 contracted gene families were majorly enriched in protein serine/threonine kinase activity, floral organ senescence, and secondary metabolite biosynthetic process (**Supplementary Figures 5-6**). In comparison with other species, 537 unique gene families were identified (**Figure 2b**) within the *S. samarangense* genome. These gene families were mainly enriched in a series of functional items, including catalytic activity, acting on DNA, retrotransfer, nucleocapsid, transfer, and RNA mediated (**Supplementary Figure 7**).

**Figure 2.**
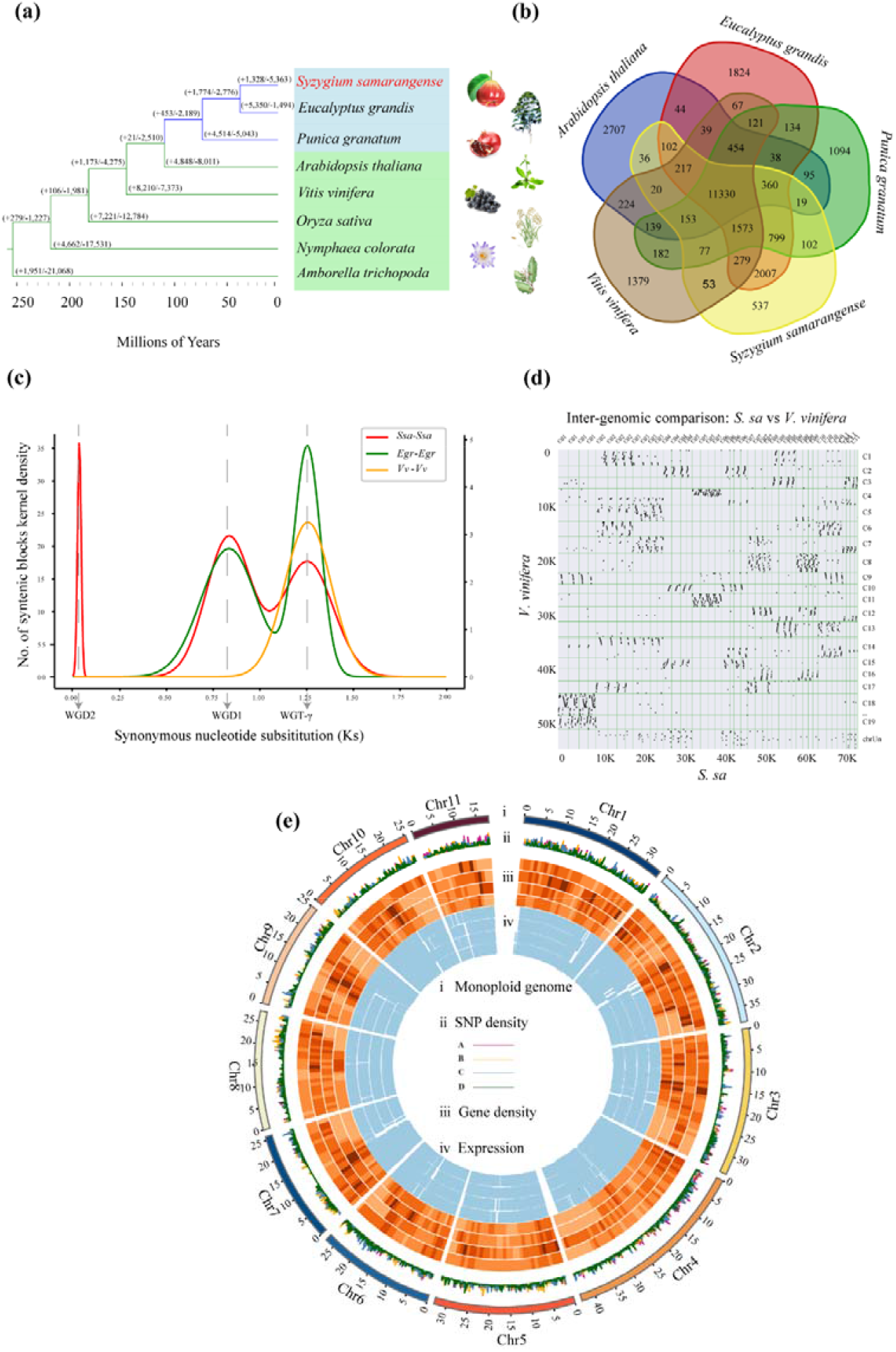
Phylogenetic and Comparative Analysis of *S. samarangense*. (a) Phylogenetic tree of *S. samarangense, E. grandis, P. granatum, A. thaliana, V. vinifera, O. sative, N. colorata*, and *A. trichopoda*. Gene family expansion/contraction analysis of the *S. samarangense* genome. The divergence times of *S. sam*arangense and the other species are labeled in the bottom. (b) Orthologous and species-specific gene families in *S. samarangense* and the other species. (c) The distribution of synonymous substitution rates (Ks) of the *S. samarangense* paralogs and orthologs with other species. (d) Alignment of *S. samarangense* genome with Vitis vinifera genome. (e) From outermost to innermost layer, these rings indicate monoploid genome in Mbp (a), SNP density among haplotypes (b), gene density (c) and expression (d), respectively. A, B, C and D respectively represents for four haplotypes in ring b, and these four haplotypes were ordered from outside to inside in rings c and d.

Comparison among the four haplotypes uncovered 4.53 million SNPs, 0.49 million short indels, and 10,925 structural variations (SVs), and these genetic variations were evenly distributed along the 44 chromosomes (**Figure 2e and Supplementary Table 8**). The clustering of chromosome-specific 13-mers partitioned each set of four haplotypes together (**Supplementary Figure 8**), which was inconsistent with the allotetraploid *Miscanthus* genome and showing the separated distribution of subgenomes. The smudge plot analysis identified that the AAAB pattern was the dominant component, accounting for 56% of examined *K*-mers (**Supplementary Figure 9**). These results collectively support that *S. samarangense* is an auto-tetraploid genome with a high level of heterozygosity.

The distribution of synonymous substitution per synonymous site (*K*_s_) of the homologous gene pairs clearly illustrated that the genome of *S. samarangense* had experienced three different rounds (WGT-γ, WGD-1, and WGD-2) of whole-genome duplication events (**Fig. 2c**). In addition to, the WGT-γ that was commonly found in the evolutionary process of grape and *E. grandis*, we discovered that *S. samarangense* and *E. grandis* had also undergone an independent whole-genome duplication (WGD-1). Compared with *E. grandis*, the specific WGD-1 event that appeared in the genome of *S. samarangense* was more complex. Moreover, the synteny relationship between the *S. samarangense* and *V. vinifera* was further analyzed to verify that WGD-1 and WGD-2 occurred after WGT-γ. As shown in **Figure 2d**, the collinear relationship between *S. samarangense* and *V. vinifera* is 8:1, indicated that the occurrence of the two lineage-specific WGDs in *S. samarangense*.

### 2.3 Genetic variations and population structure

We re-sequenced 35 accessions of *S. samarangense* at the whole-genome level and identified 2,891,846 variants, including 2,630,417 SNPs and 261,429 indels (**Supplementary Table 9**). A total of 67,430 synonymous and 78,424 non-synonymous were identified (**Supplementary Table 10**). Phylogenetic analysis demonstrated that these *S. samarangense* were partitioned into two distinct groups. The commercially cultivated accessions were clustered together as the first group, and the remaining were landraces with limited artificial selection as the second group (**Figure 3a, Supplementary Table 11**). Both principal component analysis (PCA) and genome structure were consistent with phylogenetic analysis (**Figure 3b and c**).

**Figure 3.**
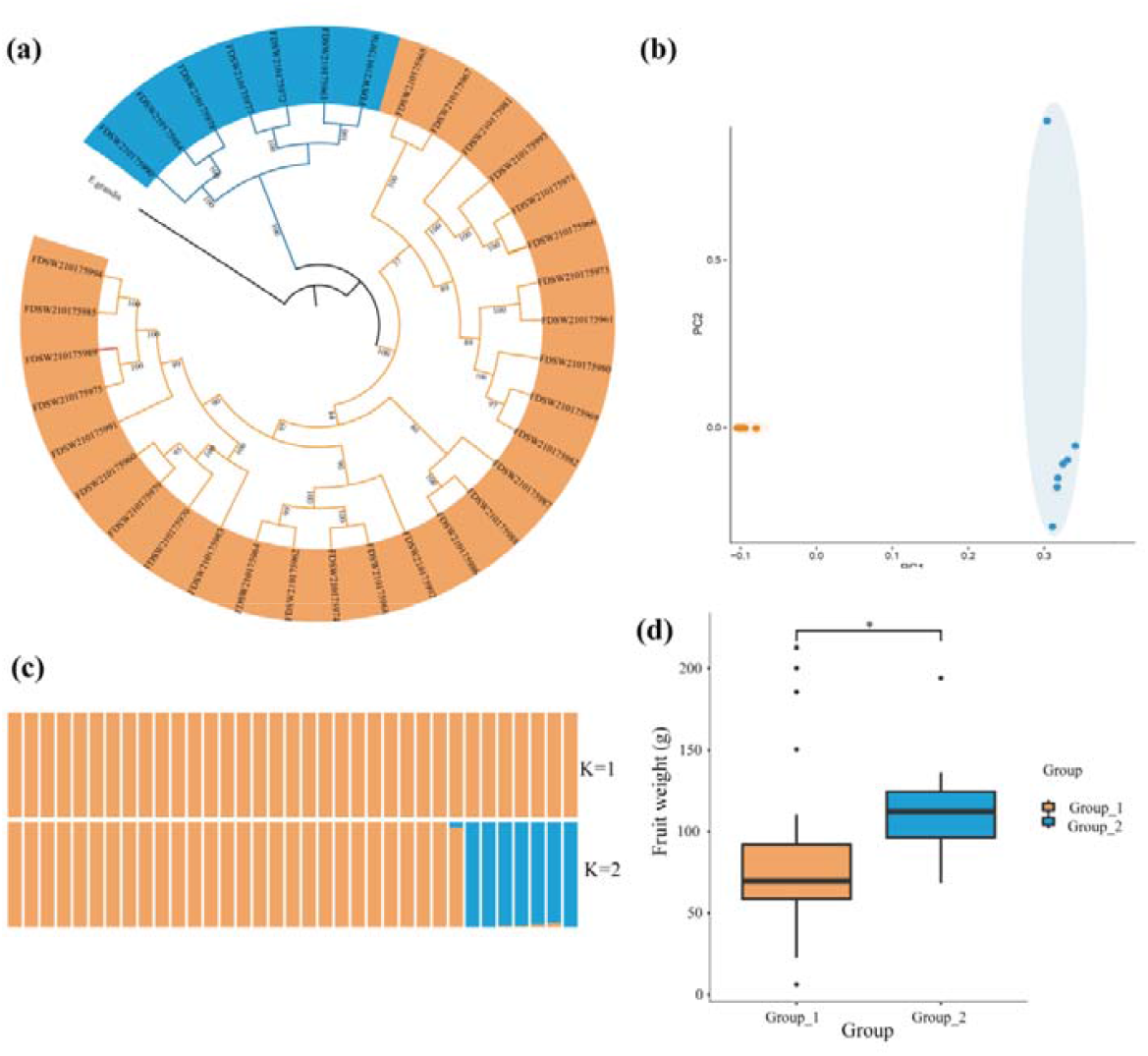
Phylogenetic splits and population genetic structure of 35 *S. samarangense* accessions. (a) Maximum-likehood tree of 35 re-sequenced *S. samarangense* individuals constructed based on 2,630,417 SNPs. (b) PCA plots of *S. samarangense* accessions showed two subgroups which indicated by different colors (blue, cultivars; yellow, landraces). PC, principle component. (c)ADMIXTRUE analysis among the accessions revealed the distribution of K=2 genetic clusters with the smallest cross-validation error. (d) Comparison of fruit weight between landraces and cultivars.

To identify the candidate genes that might have undergone natural or artificial selection during the evolutionary history in wax apple, we analyzed selective sweeps based SweeD analysis^19^ in the 35 re-sequenced individuals. A total of 22.0 Mb of genomic sequences, covering 1,299 and 1,109 protein-coding genes, were selectively swept in the landraces and cultivars, respectively. These selectively swept regions were distributed along the 11 representative chromosomes that were selected from each set of homologous chromosomes, with some chromosomes having a higher density (**Supplementary Figure 10-11**). GO enrichment analysis revealed that these swept genes were significantly enriched in the second-messenger-mediated signaling and calcium-mediated signaling pathways in landraces. However, these swept genes were enriched in metabolic process and zygote asymmetric cell division in cultivars (**Supplementary Figure 12-13**).

Phenotypic analysis showed that the cultivated wax apples had increased in fruit weight than the landraces, leading to a hypothesis that fruit growth-related genes are likely under artificial selection (**Figure 3d**). To verify this, we collected 30 homologous genes related to fruit growth in wax apple (**Supplementary table 12**) based on the published genes in tomato^20^. We observed that the landraces contained three genes located in the selectively swept genomic regions, namely *KRP1-like, IAA17-like*, and *GME-like* which has been demonstrated that involved in cell expansion, including endocycle control, auxin signaling, and ascorbate biosynthesis. In addition, the *FLACCA-like* gene which involved in ABA biosynthesis was under selection in cultivars^20^ (**Supplementary Figure 14-15**).

### 2.4 Genes contributing to fruit size and sugar content

The ‘Tub’ variety had the largest fruit weight, with an average of 124.6 g per single fruit. It is almost two times than that in ‘ZY’ (68.5 g on average) and four times in ‘DK’ (35.4 g on average). This indicated that the fruit sizes of the three varieties were significantly different. Previous study indicated that sepal development gene *APETALA* (*AP*) control the fruit size in apples^21^, which have the same fruit structure with wax apple. Through the comparative RNA-Seq data, we found that the expression of *AP1* and *AP2* genes were the highest in ‘Tub’ accession that had the largest fruit weight, followed by ‘ZY’ and ‘DK’ accessions with much reduced fruit size (**Figure 4, Supplementary Figure 16-18**). *AP1* gene was highly expressed in ‘Tub’Fr_T1 and ‘Tub’Fr_T3 samples, suggesting that *AP1* may play a role in promoting fruit growth at the early stage of fruit development.

**Figure 4.**
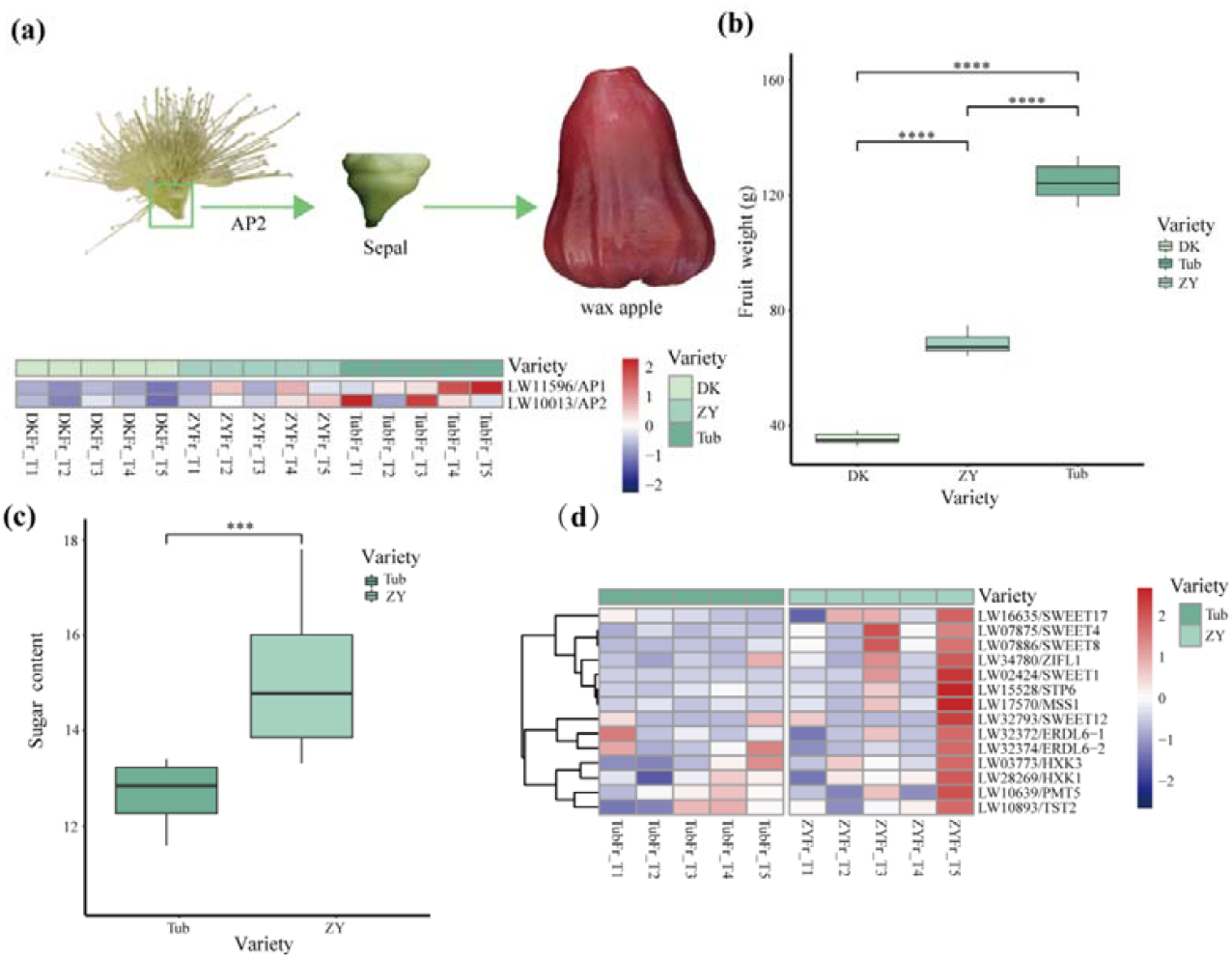
Genes related to fruit growth and sugar content. (a) The expression of sepal development homologies (*AP1* and *AP2*) in ‘DK’, ‘ZY’, and ‘Tub’ during fruit development. (b) Comparison of fruit weight among ‘DK’, ‘ZY’, and ‘Tub’. ****, *P* value < 0.0001, t-test, n = 10. (c) Comparison of sugar content between ‘Tub’ and ‘ZY’ fruit at mature. ***, *P* value < 0.001, t-test. (d) The expression of the candidate genes related to sugar transport (*SWEETs, ERDLs*, and *TST*) of pink module in ‘DK’ and ‘Tub’ during fruit development. ‘DK’: ‘Dongkeng’; ‘Tub’: ‘Tub Ting Jiang’. FrT1, FrT2, FrT3, FrT4, and FrT5 represent 10 to 50 DAFB (days after full bloom) at approximately 10-day intervals.

In fruits, sugar content is usually defined as the total soluble solid content that determines the sweetness and is an important index to determine the fruit quality. We observed that the fruits in ‘ZY’ contain a significantly higher soluble solid content than those in ‘Tub’ (14.56% v.s. 11.81%; **Figure 4c**). We further queried the meaningful genes contributing to the elevated sugar content through a comparative RNA-seq analysis of fruit samples between ‘ZY’ and ‘Tub’. WGCNA identified 14 co-expressed modules (**Supplementary Figure 19**), and a total of 400 genes were co-expressed in ‘ZY’Fr_T5 sample that is the most mature stage of ‘ZY’ fruit and assumably contains the highest level of sugar content (**Supplementary Figure 20**). We observed that a list of important sugar transporter genes exhibited significantly high levels of expression in ‘ZY’Fr_T5 (**Figure 4d**).

### 2.5 Genes associated with male sterility

Seedless fruits are highly desirable due to their commercial values. This trait is likely resulted from abnormal development of ovule and pollen^22^. The results showed that the anther contains abundant pollens and normal dehiscence in ‘DK’, but the anther contains a small amount of pollen and abnormal dehiscence in ‘Tub’ (**Figure 5a-b**). Subsequently, we detected the pollen germination rate in ‘DK’, ‘ZY’, and ‘Tub’. The results showed that the pollen germination rate was 11.73% and 45.06% for ‘ZY’ and ‘DK’ respectively, but the pollen of ‘Tub’ wasn’t collected because the anthers abnormal dehiscence (**Figure 5c**).

**Figure 5.**
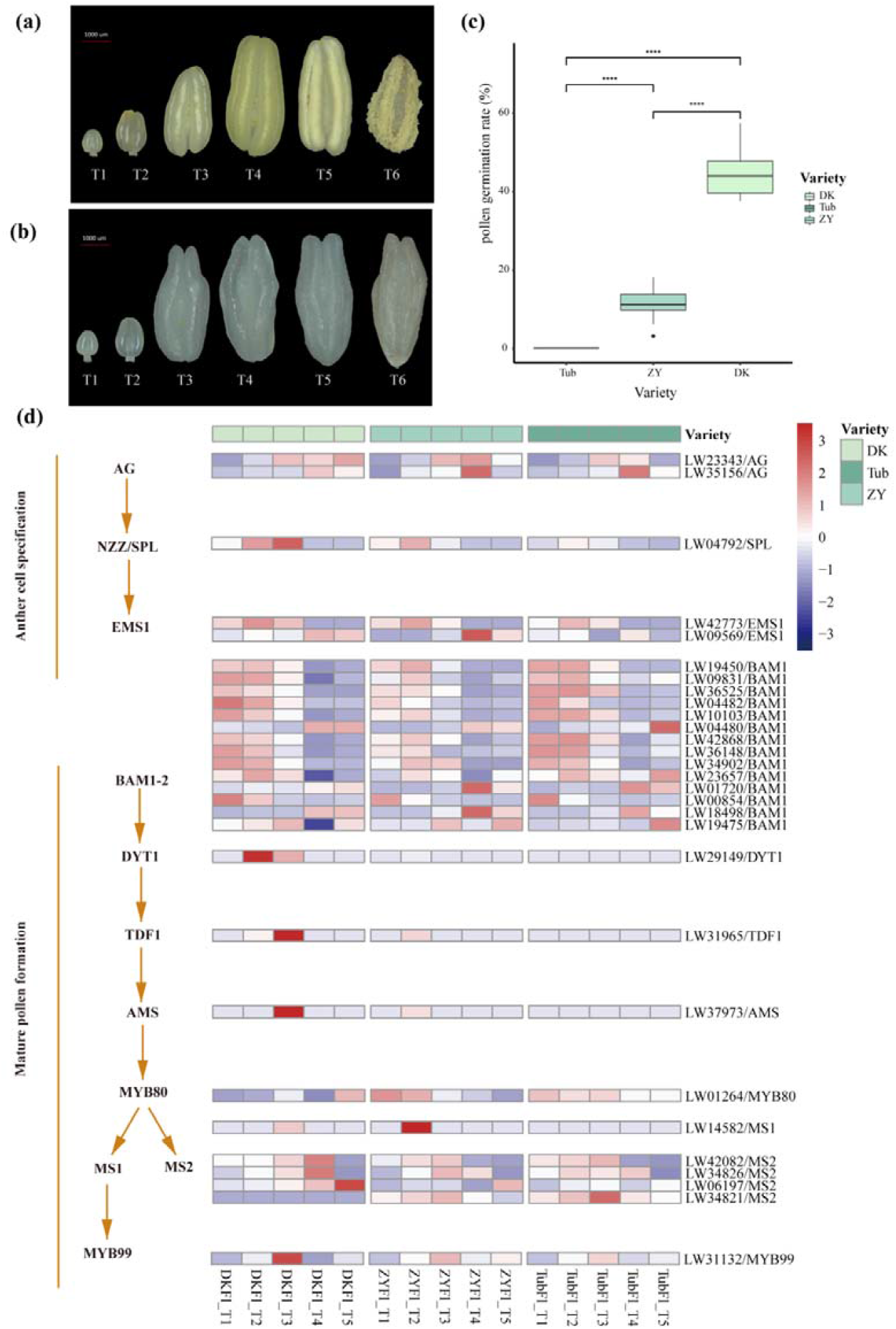
Anther development, pollen germination rate, and the expression of anther and pollen development related genes in ‘DK’, ‘ZY’, and ‘Tub’. (a) Anther development and dehiscence in ‘DK’. T1-T5 is consistent with FlT1-FlT5, and T6 represents 12 hours after blooming. (b) Anther development in ‘Tub’. T1-T5 is consistent with FlT1-FlT5, and T6 represents 12 hours after blooming. (c) Pollen germination rate of ‘Tub’, ‘ZY’, and ‘DK’. ****, *P* value < 0.0001, t-test, n = 10. (d) Expression (FPKM) of anther and pollen development related genes in ‘DK’, ‘ZY’, and ‘Tub’ from flower at different stages, including FlT1, FlT2, FlT3, FlT4, and FlT5. The expression from low to high is indicated by the scale ranging from blue to red.

In our study, the samples of different flowering stages were applied for the further RNA-seq analysis to identify key genes that involved in the development of pollens and anther. The WGCNA was performed to explore the potential genes that related to the male sterility in ‘Tub’. The coexpression network was constructed based on the correlation of gene expressions in all samples. Finally, 16 different modules, defined as the highly interconnected gene clusters, were identified and marked with different colors (**Supplementary Figure 21**). Among these modules, three potential pollen and anther development-associated module eigengenes were characterized (**Supplementary Table 13**). In ‘DK’, the turquoise, tan, and darkgreen modules were correlated with the development of pollen and anther (**Supplementary Figure 22-24**). Interestingly, the turquoise module contains the highly connected hub genes, including *LBD10, RPG1, RBOHE, CALS5, SK32*, and *MYB33*, which are known genes involved in the pollen development (**Supplementary Table 14** and **Supplementary Figure 25**).

Furthermore, we identified a total of 29 homologous genes that played an important role in male sterility in *Arabidopsis*. These genes were mainly involved in anther cell specification and mature pollen formation pathways, and many of them showed differential expression at five different flower developmental stages (FlT1 to FlT5) among the three examined varieties (**Figure 5** and **Supplementary Figure 26**). An anther cell specification related gene nozzle/sporocyteless (*NZZ/SPL*) was found to be more expressed in ‘DK’ than in ‘Tub’. We also observed that dysfunctional tapetum 1 (*DYT1*), tapetum development and function 1 (*TDF1*), and abortive microspore (*AMS*) genes were specifically expressed in FlT2 and FlT3 stages in ‘DK’, and barely expressed in ‘Tub’. In addition, the expression of three male sterile 2 (*MS2*) homologous genes in ‘DK’ was much higher than that in ‘Tub’ at FlT4 and FlT5 stages. The expression pattern of these pollen development related genes was consistent with the results of pollen germination rate (**Figure 5d**).

## 3. Discussion

The wax apple is an economically important fruit crop and widely cultivated throughout the southeast Asian countries. Here, we generated a high-quality fully phased auto-tetraploid genome assembly and 35 re-sequencing accessions. These data represented comprehensive genomic resources of this species, facilitating to investigate meaningful genetic variations and the evolutionary history. Comparative genomics and transcriptome analysis also uncovered key genes underlying fruit growth, fruit size, and sugar content, as well as factors related to male sterility caused by aborted pollen.

The assembly of wax apple is severely hindered by the high level of repetitive sequences and polyploidy. So far, only a few autotetraploid genomes were assembled to the chromosome level, including the sugarcane *Saccharum spontaneum*^18^, the cultivated alfalfa^23^, the potato cultivar^24^, and *Rehmannia glutinosa*^25^. Among these, only the sugarcane and cultivated alfalfa were assembled by combining the developed sequencing technologies and chromosome phasing algorithm, whereas the developed sequencing technologies and pollen genome were used in the potato cultivar. Here, we generated a haplotype-resolved chromosome-level genome of *S. samarangense* consisting of 44 allelic chromosomes by combining the sequencing technologies and chromosome phasing algorithm. The high percentage of assembled genome size to the monoploid estimation and anchor rate indicated a high-quality, allele-ware, and chromosome-scale genome assembly, benefiting for the downstream analysis and molecular breeding.

Fruit size and sugar content affect consumer preference. Emerging evidence shows that floral organ development related genes participate in fruit development and play different roles among species, mainly depending on the type of floral organ that develops into the fruit tissues^26^. Previous studies have shown that *AP2* governs seed yield^27^ and floral development, especially sepal development^28,29^ in *Arabidopsis* and can affect the fruit growth in apples^21^. Intriguingly, *AP2* inhibits the fruit size in *Arabidopsis*, yet promotes the fruit size in apple^21^. In apple, miR172 inhibits the expression of *AP2*, and overexpression of miR172 reduced fruit size which indicated miR172 plays a vital role in fruit size via *AP2*^21^. The high expressions of sepal development genes (*AP1* and *AP2*) in our results were in the ‘Tub’ group that had the greatest fruit weight, which suggests that *AP1*/*2* may play an important role in the regulation of wax apple fruit size. Considering that the wax apple is recognized as the false fruit which develops from the ovary and sepals, the genes regulating sepal development were likely related to fruit size. *APETALA2* (*AP2*) governs sepal development, and *APETALA2* (*AP1*) acts downstream of *AP2*^30^. The main reason for the phenomenon is that unlike the fruits of apple and *S. samarangense* that grow from the sepals, the siliques of *Arabidopsis* develop from ovary tissues^31^. In apple, overexpression of *MdERDL6*-1 improved the glucose (Glc), fructose (Fru), and sucrose (Suc) concentration in transgenic apple fruit and increased the expression of *TST1*/*TST2* indicating that the sugar content in vacuoles were mediated by the co-ordinated action of *MdERDL6*-1 and *MdTST1/2*^32^. In our study, *ERDL6*-1 and *TST2* were mainly expressed in ‘ZY’ variety which contains higher sugar content, indicating that the sugar accumulation in ‘ZY’ variety is possibly attributed to the higher expression of *ERDL6*-1 and *TST2*. Through a comparative RNA-seq analysis of fruit samples for the meaningful genes contributing to the elevated sugar content between ‘ZY’ and ‘Tub’, we identified 14 co-expressed modules, and a total of 400 genes were co-expressed in ‘ZY’Fr_T5 sample that is the most mature stage of ‘ZY’ fruit and assumably contains the highest level of sugar content, these genes were significantly enriched in a series of molecular functions, particular in sugar transporter activity items. In addition, the high levels of expression in ‘ZY’Fr_T5 for list of important sugar transporter genes including sucrose transporters (*SUTs*), monosaccharide transporters (*MSTs*), and sugars will eventually be exported transporters (*SWEETs*) and *TMT2*. Our results collectively supported that these sugar transporter-related genes contributed to elevated sugar content in the fruit of wax apple.

Seedless fruit occupies an important position in the domestic and international market. in *Arabidopsis* the *LBD10* ortholog can interact with *LBD27* to form a heterodimer and plays an essential role in the pollen development^33^, highly suggesting its potential role in the regulation of male sterility in wax apple, and many species have been developed, such as grape and Citrus^34,35^. Male sterility caused by aborted pollen is the main pathway to cultivate seedless fruit. Based on these evidences, we speculate that the male sterility in ‘Tub’ is possibly attributed to functional defects of a couple of key genes, especially *DYT1, TDF1*, and *AMS*, affecting the early tapetum development. In *Arabidopsis*, previous investigations showed that *DYT1, TDF1, AMS* mutants all display a fully male sterile phenotype^36-38^. *DYT1-TDF1-AMS-MS188* genetic network was suppressed in the mutation of *Fatty Acid Export 1* and caused defective pollen formation^39^. Trace concentrations of imazethapyr (IM) results the gene expression of *DYT1, TDF1*, and *AMS* decreased significantly, which affected anther and pollen biosynthesis in *Arabidopsis*^40^. Here, we identified that *DYT1, TDF1*, and *AMS* were highly expressed in male fertile variety ‘DK’, but lower in ‘Tub’, and finally in male sterile variety ‘ZY’. Therefore, those genes may play the potential role in the regulation of fertility in wax apple. Together, male sterility produces seedless fruit and may be caused by the decreased expression of *DYT1, TDF1*, and *AMS*. The results suggested that these genes could play important roles in the seedless phenotype formation, and the relative expression level in *LBD10, RPG1, RBOHE, CALS5, SK32*, and *MYB33* versus *DYT1, TDF1*, and *AMS* seemed to be key factor in this process in wax apple.

## 4. Conclusions

Here, a haplotype-resolved autotetraploid genome assembly of the wax apple was generated, and comparative genomic analysis revealed *S. samarangense* had experienced three different rounds of WGD events, including two independent WGDs after WGT-γ. Transcriptome analysis was used to identify the genes related to fruit size, sugar content, and male sterility. Combined with fruit weight, fruit development characteristics, and transcriptome data analysis, *AP1* and *AP2* genes may regulate fruit size by regulating sepal development. Sugar transport-related genes (*SWEETs* and *SUTs*) was found to be higher expressed in variety with higher sugar content in ‘ZY’. The low expression of *DYT1, TDF1*, and *AMS* in ‘Tub’ may be the main reason for its sterility. Our results provide the foundation for further study on the regulatory mechanisms of fruit quality and male sterility, and can be used in molecular assisted breeding of wax apple, especially for seedless traits.

## 5. Methods

### 5.1 Illumina short-read sequencing and genome survey

We chose the ‘Tub’ accession for *de novo* genome sequencing and assembly. The plant materials were maintained by Fujian Academy of Agricultural Sciences, and young leaves were collected from an individual tree planted in the Field GenBank for wax apple of Fujian Academy of Agricultural Sciences, Fujian province, China (Coordinates: 26°7′53″N; 119°20′6″E) under the voucher number GPLWFJGSS0058. Genomic DNA was isolated from young leaves using the Qiagen Plant Genomic DNA Kit according to the manufacturer’s instructions. Then, the qualified DNA samples were randomly fragmented with a Covaris S-series Instrument, and Illumina PCR-free libraries with insert sizes of 350-bp were constructed using Truseq Nano DNA HT Sample preparation Kit (Illumina USA). Finally, the constructed libraries were sequenced with 150-bp paired-end sequencing using Illumina HiSeq PE. Using Illumina short reads, the genome size, repeat contents, and heterozygosity rate of *S. samarangense* were estimated using jellyfish2.2.7 software^41^.

### 5.2 Genome sequencing

A combination of single-molecule real-time sequencing (SMRT), Illumina sequencing, and Hi-C sequencing with error correction was applied to assemble the complete genome sequence of *S. samarangense*. For SMRT, genomic DNA was disrupted randomly with 6 kb-20 kb fragments by g-TUBE (Covaris, Woburn, MA, USA) and sequenced by the PacBio Sequel platform, generating 110 coverage. For Illumina sequencing, 6 libraries (300 bp) were constructed using Illumina Truseq Nano DNA Library Prep kit, and the libraries were sequenced on the Illumina Hi-Seq 2000 platform. For Hi-C sequencing, two Hi-C libraries were constructed using a standard procedure and sequenced using the Illumina Hiseq X Ten sequencer.

### 5.3 Genome assembly

The contig-level assembly of the wax apple genome incorporated Illumina short reads and PacBio CLR subreads. The PacBio subreads were subject to the whole pipeline of Canu assembler v1.9^16^, followed by the polishing using the Pilon program^42^ to increase assembly accuracy. To construct the haplotype-resolved genome assembly, we first mapped the Hi-C reads to the polished contigs assembly using BWA MEM (−5SPM) and extract the uniquely mapped paired reads. The resulting BAM files were applied on haplotype phasing and scaffolding using ALLHiC pipeline^43^. In addition, the chimeric scaffolds were manually corrected based on the Hi-C signals in juicebox. To fill the gaps, first, TGS GapCloser^44^ software was used to fill the gaps in the wax apple genome with 30X ultra-long ONT data. After filling the genome, the number of gaps were significantly reduced. Then, we used Merqury^45^ software to check the gap filled genome, and found that some errors were introduced compared with the previous filling. To correct these errors, we extracted all gap sequences filled by TGS GapCloser, and checked the QV quality value of each gap and the error rate of the corresponding sequence in the genome using the Mercury software. Finally, we filled the correct GAP into the initial chromosomal level genome. The quality of chromosome-scale assembly was assessed using Hi-C heatmap.

### 5.4 Genome annotation

To annotate protein-coding genes, we followed the method described in the previous study^46^. Briefly, we integrated evidences from RNA-seq, orthologous proteins, and ab initio gene prediction by carrying out two rounds of MAKER pipeline. In the first round of MAKER, Trinity was used to de novo assembly by using the RNA-seq data^47^ and RSEM was applied to calculate transcript abundance^48^. After filtering the valid transcript, the rest were imported to the PASA program and the candidate proteins were trained by the ab initio gene prediction^49^. In the second round of MAKER, the candidate proteins were retrained by ab initio. Hisat2 and StringTie were used to reassemble^50,51^. Finally, we selected the better annotation of the two rounds annotation. The BUSCO (v.5) software was applied to calculate the degree of annotation complement. We used the same method as describing in an autopolyploid sugarcane genome to construct a monoploid genome, identify alleles, and analyze allelic variations^18^.

### 5.5 RNA library construction and sequencing

To improve the prediction of gene annotation, we performed RNA-seq using different tissues of *S. samarangense* including flesh, flower, leaf, ovary, root, and stem. All these tissues were collected and subsequently frozen in liquid nitrogen. Total RNA was extracted with the RNAprep Pure Plant Plus Kit (TIANGEN) following the manufacturer’s procedure. Transcriptome libraries were constructed using NEBNext® Ultra™ RNA Library Prep Kit for Illumina (NEB, UK) according to the manufacturer’s instructions and sequenced with 150-bp paired-end sequencing using the Illumina NovaSeq 6000 (Illumina, USA) platform.

### 5.6 Phylogenetic analysis and estimation of the divergence time

To construct the phylogenetic tree, single-copy orthologous genes were defined by OrthoFinder v2.3.1^52^ from protein sequences of seven species (*Eucalyptus grandis, Punica granatum, Arabidopsis thaliana, Vitis vinifera, Oryza sativa, Nymphaea colorata*, and *Amborella trichopoda*). Afterwards, protein sequences were aligned by MUSCLE^53^ and GBLOCKS^54^ was used to trim ambiguous alignment portions. A phylogenetic tree was constructed using RAxML^55^ utilizing the JTT+I+G+F model and 1,000 bootstrap analyses. The divergence time among these species was estimated by r8s^56^. Whether the gene families had undergone the expansion or contraction events in the eight sequenced species were identified using CAFE2.2^57^.

### 5.7 Synteny and whole-genome duplication analysis

To investigate the whole genome duplication (WGD) events in *S. samarangense*, synteny analysis of *S. samarangense* and *V. vinifera* genome was performed. The *V. vinifera* genome and annotation were downloaded from phytozome (https://phytozome-next.jgi.doe.gov/). We applied the MCScan (python version) pipeline^58^ following the suggested best workflow. The syntenic regions in *S. samarangense* and *V. vinifera* genome supported that *S. samarangense* experienced two WGD events after WGT-γ.

To test the reliability of this result, the synonymous nucleotide substitutions on synonymous sites (Ks) values in *S. samarangense, V. vinifera*, and *E. grandis* genomes were estimated by YN00 program in the WGDi package with the Nei-Gojobori approach^59^. For the base substitution rate is different in the three species, the method applied by Jinpeng Wang^60^ was used to correct the evolutionary rate of duplicated genes. After fit and merge operations, the Ks peaks caused by the same WGD event could locate in the same place.

### 5.8 Resequencing and population analysis

A total of 35 accessions were re-sequenced, including 28 landraces and seven cultivars. All accessions were collected from the Field GenBank for wax apple of Fujian Academy of Agricultural Sciences, Fujian province, China. Young leaves were collected from each accession and flash frozen in liquid nitrogen for DNA isolation. Genomic DNA from each sample was isolated, and paired-end reads were sequenced on the Illumina NovaSeq platform. The adaptors and low-quality were trimmed using Trimmomatic^61^, and clean reads were aligned to the reference genome of *S. samarangense* using BWA with default parameters^62^. We identified variants following the GATK^63^ best practices pipeline. HaplotypeCaller and GenotypeCaller were used to call variants from all samples. Maximum-likelihood trees were constructed using VCF2Dis (https://github.com/BGI-shenzhen/VCF2Dis).

To infer the subgroup among the re-sequenced *S. samarangense* accessions, admixture^64^ was used with different *k* values (from 1 to 3), the optimal value determined in this study was *k*=2. PLINK1.9, and VCFtools^65^ were used to perform PCA. Finally, we used SweeD^19^ to detect complete selective sweeps in the *S. samarangense* genome with default settings.

### 5.9 Transcriptome sequencing and identification of co-expression modules

The fruits from three wax apple accessions, ‘ZY’, ‘Tub’, and ‘DK’, were sampled from 10 to 50 DAFB (days after full bloom) at approximately 10-day intervals, representing five developmental stages, namely T1 to T5. Total RNA was extracted from flower and fruit using RNAprep Pure Plant Plus Kit (TIANGEN), cDNA libraries were constructed and sequenced by Illumina NovaSeq 6000 (Illumina, USA) platform. Subsequently, we evaluated reads quality by FastQC software (http://www.bioinformatics.babraham.ac.uk/projects/fastqc), removed sequencing adapters and low-quality bases using Trimmomatic^61^. The clean data were aligned to the *S. samarangense* genome using HISAT2 (v2.0.5)^66^, and the fragments per kilobase per million mapped fragments (FPKM) value was calculated using StringTie (v.1.2.3)^67^. The R package weighted gene co-expression network analysis (WGCNA) was used to cluster genes with similar expression based on the FPKM data^68^. Genes with |MM|>0.8 and |GS|>0.2 were selected for further analysis, and the network was represented and displayed using Cytoscape (v.3.6.0)^69^. Male sterility and flower development related genes were retrieved from *Arabidopsis* (https://www.arabidopsis.org/), and the homologs of *S. samarangense* were identified by BLASTP search of these sequences against all *S. samarangense* protein sequences.

### 5.10 Fruit quality analysis and pollen viability determination

For fruit weight analysis, the fruits of all 35 *S. samarangense* materials (including 28 landraces and seven cultivars) were collected. Ten fruits were randomly selected from three trees for each *S. samarangense* material. The fruit weight was measured by electronic balance QUINTIX213-1CN (Sartorius, Germany). To determine the total soluble solids (TSS) content, take 1 cm^3^ of tissue from the upper, middle, and lower parts of the each fruit sample, respectively. Then mixed and homogenized them thoroughly with a mortar and pestle. The supernatant of the homogenate was used for soluble solids content determinations by a hand-held Brix meter PAL-1 (ATAGO, Japan). To analyze pollen viability, pollen tube germination rate was measured. At 35 °C, the pollen was cultured for 12 hours in the medium (the concentration of sucrose was 15%, the concentration of boric acid was 50mg/L, and the concentration of agar was 1%). Then, optical microscope was used to observe the pollen tube germination. Three fields of vision were randomly selected, the total number of pollen and the number of germinated pollens were counted at the same time. The germination rate was calculated. The standard for budding pollen is: the length of the pollen tube exceeds the diameter of the pollen.

For each experiment, the significance of between-group differences was analyzed using t-test. All statistical analyses were performed using IBM SPSS software. *p*-value < 0.001 was considered to be statistically significant.

## Acknowledgements

This work was supported by the Natural Science Foundation of Fujian Province (2020J011361), the High-quality Development beyond the ‘ 5511 ’ Collaborative Innovation Project in Fujian province (XTCXGC2021016-4). We thank Ping Zhou (Fujian Academy of Agricultural Sciences) for analysis program in data.

## Data availability

Raw sequencing reads used for de novo whole-genome assembly and the final genome have been deposited in the WGS under access number WGHBKKI00000000. Raw resequencing data were uploaded to National Genomics Data Center (NGDC, https://ngdc.cncb.ac.cn/), submission ID: WGS034963; BioProject access number: PRJCA011822; Biosample access number: SAMC1129200; GSA access number: CRA010157.

## Author contributions

Jiahui Xu and Lihui Zeng designed the experiments; Xiuqing Wei performed the most of the experiments; Min Chen performed the genome assembly, annotation and the transcriptome data analysis; Xijuan Zhang and Lin Xu performed phenotype analysis; Liang Li collected the materials for sequencing; Huanhuan Wang and Caihui Wang analyzed the resequenced data; Mengwei Jiang conducted comparative genomic analysis. Yinghao Wang filled the gaps of wax apple genome.

