## Supplemental Table 1-12,14 and Supplementary Figure 1-26 for "The Haplotype-resolved Autotetraploid Genome Assembly Provides Insights into the genomic evolution and fruit divergence in Wax apple (*Syzygium samarangense* (BI.) Merr.et Perry)"

**Supplementary Table 1	Summary of PacBio long reads.**

| PacBio Sequel Ⅱ sequencing | *Syzygium samarangense* |
| --- | --- |
| Raw data (Gb) | 92.0 |
| Sequencing depth (x) | 110 |
| Average reads length (bp) | 11,722 |
| Reads N50 (bp) | 15,054 |
| Illumina short reads | 90.0 |
| Raw data (Gb) | 90.0Gb |
| Hi-C reads | 92.4 |

**Supplementary Table 2 Statistics of pre-assembly of the *Syzygium samarangense* genome**

| Items | assembly |
| --- | --- |
| Assemble size (Mbp) | 1,485 |
| No. of contigs | 25,179 |
| Maximum length (bp) | 4,225,000 |
| N90(bp) | 60,278 |
| N80(bp) | 75,000 |
| N70(bp) | 95,147 |
| N60(bp) | 195,107 |
| N50(bp) | 304,482 |
| Average length(bp) | 148,188 |

**Supplementary Table 3. BUSCO assessment of genome assemblies.**

| **Description** | **Number** | **Percentage (%)** |
| --- | --- | --- |
| Complete BUSCOs (C) | 1597 | 98.9 |
| Complete and single-copy BUSCOs (S) | 233 | 14.4 |
| Complete and duplicated BUSCOs (D) | 1364 | 84.5 |
| Fragmented BUSCOs (F) | 11 | 0.7 |
| Missing BUSCOs (M) | 6 | 0.4 |
| Total BUSCO groups searched | 1614 | 100 |

**Supplementary Table 4. Assessment of genome consistency**

| **Iterms** | **Statistics** |
| --- | --- |
| Number of reads | 600,624,892 |
| Data size (Gb) | 90 |
| Mapped bases (Gb) | 86 |
| Map rate (%) | 95.6 |
| Mean Depth | 75.7 |
| Coverage Rate (%) | 97.9 |

**Supplementary Table 5. Annotation completeness assessment of the *Syzygium samarangense* genome using BUSCO.**

| **Description** | **Number** | **Percentage (%)** |
| --- | --- | --- |
| Complete BUSCOs (C) | 1464 | 90.7 |
| Complete and single-copy BUSCOs (S) | 345 | 21.4 |
| Complete and duplicated BUSCOs (D) | 1119 | 69.3 |
| Fragmented BUSCOs (F) | 89 | 5.5 |
| Missing BUSCOs (M) | 61 | 3.8 |
| Total BUSCO groups searched | 1614 | 100 |

**Supplementary Table 6 Allele annotation in** ***S. samarangense.***

|  | Total count of allele genes | No.of genes with 4 alleles | No.of genes with 3 alleles | No.of genes with 2 alleles | No.of genes with 1 alleles | No.of tandem duplicated genes | No.of dispersely duplicated genes |
| --- | --- | --- | --- | --- | --- | --- | --- |
| Chr01 | 3,048 | 247 | 1,120 | 1,059 | 622 | 101 | 1,234 |
| Chr02 | 2,567 | 338 | 850 | 879 | 500 | 129 | 1,516 |
| Chr03 | 2,703 | 210 | 708 | 908 | 877 | 94 | 1,134 |
| Chr04 | 1,907 | 284 | 626 | 667 | 330 | 116 | 1,350 |
| Chr05 | 2,060 | 322 | 580 | 812 | 346 | 86 | 1,189 |
| Chr06 | 2,242 | 137 | 690 | 876 | 539 | 96 | 902 |
| Chr07 | 2,238 | 91 | 534 | 949 | 664 | 63 | 808 |
| Chr08 | 1,858 | 199 | 585 | 659 | 415 | 77 | 995 |
| Chr09 | 2,009 | 125 | 534 | 870 | 480 | 62 | 703 |
| Chr10 | 2,140 | 123 | 660 | 874 | 483 | 79 | 769 |
| Chr11 | 1,244 | 64 | 387 | 468 | 325 | 49 | 552 |
| Gene with annotated alleles | 24,016 | 2,140 | 7,274 | 9,021 | 5,581 | - | - |
| Duplicate genes | 12,113 | - | - | - | - | 952 | 11,161 |
| Unanchored genes/alleles | 555 |  |  |  |  |  |  |

**Supplementary Table 7 Statistics of repetitive elements in *S.samarangense* Genome**

| **Class** | **Count** | **bpMasked** | **%masked** |
| --- | --- | --- | --- |
| **LTR** | 431,494 | 385,171,537 | 24.74% |
| Copia | 119,285 | 89,702,677 | 5.76% |
| Gypsy | 180,121 | 229,125,587 | 14.72% |
| unknown | 132,088 | 66,343,273 | 4.26% |
| **TIR** | 543,208 | 169,848,407 | 10.9% |
| CACTA | 111,826 | 32,921,473 | 2.11% |
| Mutator | 321,401 | 95,353,532 | 6.12% |
| PIF_Harbinger | 35,231 | 13,576,572 | 0.87% |
| Tc1_Mariner | 13,521 | 4,647,381 | 0.30% |
| hAT | 612,29 | 23,349,449 | 1.50% |
| **nonTIR** |  |  |  |
| helitron | 133,738 | 38,230,905 | 2.46% |
| **Total** | 1,108,440 | 593,250,849 | 38.10% |

**Supplementary Table 8 Characteristics of genetic variation compared with monoploid genome in *S.samarangense***

| **Haplotype** | **Variation** | **Chr1** | **Chr2** | **Chr3** | **Chr4** | **Chr5** | **Chr6** | **Chr7** | **Chr8** | **Chr9** | **Chr10** | **Chr11** |
| --- | --- | --- | --- | --- | --- | --- | --- | --- | --- | --- | --- | --- |
| **A** | SNPs | 172,089 | 189,784 | 91,872 | 128,068 | 136,787 | 125,978 | 86,829 | 105,341 | 81,124 | 88,670 | 48,791 |
|  | No. of Indels(1-10bp) | 22,922 | 26,538 | 8,113 | 12,888 | 18,232 | 17,111 | 11,367 | 13,970 | 11,504 | 12,357 | 4,817 |
|  | No. of large Indels(>10bp) | 40 | 49 | 274 | 412 | 42 | 28 | 19 | 29 | 18 | 32 | 195 |
|  | Size of Indels(1-10bp) | 24,947 | 28,886 | 15,645 | 23,633 | 20,133 | 18,757 | 12,384 | 15,318 | 12,493 | 13,409 | 9,255 |
|  | Size of Indels(>10bp) | 1,226 | 1,683 | 5,276 | 8,414 | 1,532 | 965 | 506 | 969 | 644 | 852 | 3,698 |
|  | No. of Repeat expansion/contraction | 16 | 13 | 181 | 259 | 7 | 9 | 12 | 6 | 5 | 3 | 90 |
|  | Size of Repeat expansion/contraction | 18,289 | 184,926 | 16,055,635 | 19,284,101 | 3,987 | 59,770 | 102,850 | 161,819 | 16,907 | 3,216 | 6,570,209 |
| **B** | SNPs | 153,562 | 149,578 | 109,733 | 156,699 | 128,121 | 83,122 | 88,598 | 108,433 | 67,588 | 76,117 | 58,382 |
|  | No. of Indels(1-10bp) | 14,388 | 16,564 | 13,881 | 21,325 | 12,535 | 8,208 | 8,170 | 10,002 | 6,993 | 7,132 | 5,451 |
|  | No. of large Indels(>10bp) | 500 | 522 | 17 | 60 | 349 | 263 | 304 | 382 | 226 | 229 | 205 |
|  | Size of Indels(1-10bp) | 28,102 | 29,214 | 15,155 | 23,203 | 22,940 | 14,838 | 15,887 | 19,314 | 13,078 | 13,720 | 9,729 |
|  | Size of Indels(>10bp) | 9,394 | 10,212 | 749 | 2,111 | 6,590 | 5,033 | 5,761 | 7,340 | 4,241 | 4,439 | 3,828 |
|  | No. of Repeat expansion/contraction | 253 | 13 | 3 | 12 | 208 | 144 | 172 | 249 | 139 | 131 | 102 |
|  | Size of Repeat expansion/contraction | 16,640,249 | 172,783 | 47860 | 30,893 | 15,885,447 | 7,419,693 | 8,988,695 | 10,901,617 | 8,084,148 | 10,496,674 | 7,312,797 |
| **C** | SNPs | 124,016 | 126,197 | 94,058 | 121,475 | 100,771 | 118,582 | 60,773 | 68,391 | 65,674 | 73,687 | 74,361 |
|  | No. of Indels(1-10bp) | 11,871 | 12,484 | 8,313 | 11,838 | 9,260 | 11,315 | 5,927 | 6,244 | 6,203 | 6,611 | 9,499 |
|  | No. of large Indels(>10bp) | 408 | 442 | 301 | 424 | 343 | 397 | 194 | 251 | 238 | 237 | 15 |
|  | Size of Indels(1-10bp) | 21,920 | 22,637 | 16,381 | 22,143 | 17,481 | 21,493 | 10,672 | 12,088 | 11,914 | 12,938 | 10,197 |
|  | Size of Indels(>10bp) | 7,769 | 8,664 | 5,596 | 8,357 | 6,238 | 7,427 | 3,624 | 4,701 | 4,558 | 4,701 | 516 |
|  | No. of Repeat expansion/contraction | 223 | 272 | 181 | 225 | 167 | 229 | 107 | 126 | 125 | 147 | 7 |
|  | Size of Repeat expansion/contraction | 14,185,638 | 24,902,938 | 12,861,477 | 18,038,207 | 16,681,132 | 13,373,520 | 7,182,981 | 12,093,672 | 9,246,022 | 7,496,460 | 186,881 |
| **D** | SNPs | 140,709 | 176,883 | 68,395 | 118,702 | 111,732 | 107,455 | 57,323 | 81,066 | 61,244 | 75,203 | 68,289 |
|  | No. of Indels(1-10bp) | 13,492 | 17,292 | 6,385 | 11,647 | 10,942 | 10,628 | 5,649 | 7,218 | 5,672 | 7,858 | 6,388 |
|  | No. of large Indels(>10bp) | 488 | 587 | 181 | 400 | 391 | 312 | 165 | 291 | 195 | 240 | 230 |
|  | Size of Indels(1-10bp) | 24,825 | 32,412 | 11,678 | 21,333 | 20,577 | 20,270 | 10,250 | 13,830 | 11,224 | 14,719 | 12,084 |
|  | Size of Indels(>10bp) | 9,432 | 10,967 | 3,508 | 7,774 | 7,690 | 5,912 | 3,173 | 5,355 | 3,709 | 4,520 | 4,593 |
|  | No. of Repeat expansion/contraction | 268 | 223 | 131 | 222 | 201 | 200 | 126 | 172 | 119 | 148 | 122 |
|  | Size of Repeat expansion/contraction | 16,834,811 | 25,136,188 | 7,574,176 | 24,164,469 | 13,971,299 | 11,586,184 | 8,784,594 | 13,729,699 | 4,782,982 | 10,602,422 | 10,457,504 |

| **Variation type** | **Number** |
| --- | --- |
| SNP | 2630417 |
| InDel | 261429 |
| Total | 2891846 |

**Supplementary Table 9 Statistics of variants among the 36 re-sequenced *S. samarangense*.**

**Supplementary Table 10 Statistics of variation type and region among the 36 re-sequenced *S.samarangense***

| **Variation type and region** | **Count** | **Percent** |
| --- | --- | --- |
| 3_prime_UTR_variant | 62826 | 1.223% |
| 5_prime_UTR_premature_start_codon_gain_variant | 7685 | 0.15% |
| 5_prime_UTR_truncation | 1 | 0% |
| 5_prime_UTR_variant | 50956 | 0.992% |
| Bidirectional_gene_fusion | 4 | 0% |
| conservative_inframe_deletion | 309 | 0.006% |
| conservative_inframe_insertion | 554 | 0.011% |
| disruptive_inframe_deletion | 615 | 0.012% |
| disruptive_inframe_insertion | 608 | 0.012% |
| downstream_gene_variant | 1008550 | 19.627% |
| Exon_loss_variant | 4 | 0% |
| frameshift_variant | 4234 | 0.082% |
| Gene_fusion | 1 | 0% |
| initiator_codon_variant | 13 | 0% |
| intergenic_region | 2319469 | 45.138% |
| intron_variant | 401309 | 7.81% |
| missense_variant | 76309 | 1.485% |
| Non_coding_transcript_variant | 70 | 0.001% |
| splice_acceptor_variant | 374 | 0.007% |
| splice_donor_variant | 347 | 0.007% |
| splice_region_variant | 10984 | 0.214% |
| start_lost | 184 | 0.004% |
| stop_gained | 1727 | 0.034% |
| stop_lost | 204 | 0.004% |
| stop_retained_variant | 100 | 0.002% |
| synonymous_variant | 67430 | 1.312% |
| upstream_gene_variant | 1123780 | 21.869% |

**Supplementary Table 11 Statistics of the 35 re-sequenced accessions in this study.**

| Group |  | Name of Germplasm Resources | Single Fruit Weight (g) | Places of Origin |
| --- | --- | --- | --- | --- |
| Group1 | FDSW210175985 | XiangChengBenDi | 69.06 | Fujian,China |
|  | FDSW210175994 | GuiPingBenDi | 58.62 | Guangxi,China |
|  | FDSW210175975 | NanYa 17 | 69.44 | Thailand |
|  | FDSW210175989 | DaHongZhong | 56.61 | Hainan,China |
|  | FDSW210175991 | HaiNanBenDi | 22.89 | Hainan,China |
|  | FDSW210175992 | JinZuan | 73.28 | Thailand |
|  | FDSW210175983 | FeiCui | 96.66 | Taiwan,China |
|  | FDSW210175970 | FenHong | 69.91 | Fujian,China |
|  | FDSW210175960 | HeiZuan | 83.2 | Taiwan,China |
|  | FDSW210175979 | TaiNong 2 | 93.12 | Taiwan,China |
|  | FDSW210175987 | Peth Sam Pung | 59.4 | Thailand |
|  | FDSW210175986 | Toon Klaw | 62.7 | Thailand |
|  | FDSW210175988 | MaLaiXiYaQingZhong | 110.411875 | Malaysia |
|  | FDSW210175976 | XiShuangBanNa 6 | 59.69285714 | Thailand |
|  | FDSW210175968 | ShunDeBenDi | 48.7 | Gongdong,China |
|  | FDSW210175962 | TaiGuoQingZhong | 63.36631579 | Thailand |
|  | FDSW210175964 | HeiZuanShi | 83.2 | Taiwan,China |
|  | FDSW210175967 | NongKe 4 | 200.124 | Fujian,China |
|  | FDSW210175965 | YinNiDaGuo | 212.946 | Indonesia |
|  | FDSW210175982 | HeiTangBaBi | 150.44 | Taiwan,China |
|  | FDSW210175980 | QingZuan | 89.08 | Fujian,China |
|  | FDSW210175969 | ShuangSe | 78.34 | Fujian,China |
|  | FDSW210175981 | DongKeng 3 | 35.41 | Fujian,China |
|  | FDSW210175993 | BaiLianWu | 6.171 | Yunnan,China |
|  | FDSW210175971 | YinDuHong | 185.753 | India |
|  | FDSW210175966 | LongWenBenDi | 35.69 | Fujian,China |
| Group2 | FDSW210175973 | DaYeHong | 95.66 | Indonesia |
|  | FDSW210175961 | DaYeLianWu | 122.075 | Indonesia |
|  | FDSW210175978 | MiFengLing | 96.37 | Taiwan,China |
|  | FDSW210175977 | Tub Ting Jiang | 124.6 | Thailand |
|  | FDSW210175972 | HeiJinGang | 102.95 | Taiwan,China |
|  | FDSW210175978 | ZiYu | 68.47 | Fujian,China |
|  | FDSW210175984 | BaZhangLianWu | 194.164 | Taiwan,China |
|  | FDSW210175990 | FeiDan | 112.264 | Hainan,China |
|  | FDSW210175963 | TaiGuoHongBaoShi | 136.351 | Thailand |

**Supplementary Table 12 Fruit growth-related genes in *S. samarangense*.**

| Ovary Dev | Carpel identity and Cell division control | TAGL1 | LW35156 |  |  |
| --- | --- | --- | --- | --- | --- |
|  |  | FAS | LW06224 |  |  |
|  |  | LC | LW08101 |  |  |
| Fruit set | Auxin signaling | SlPIN4 | LW14700 |  |  |
|  |  | SlTIR1 | LW33579 |  |  |
|  |  | SlARF7 | LW28041 |  |  |
|  |  | SlARF8 | LW29664 |  |  |
|  |  | SlIAA9 | LW02730 |  |  |
|  | GA signaling | SlGA20ox1 | LW36126 |  |  |
|  |  | SlDELLA1 | LW38360 | LW07312 |  |
| Cell division | Cell division control | FW2.2 | LW10547 |  |  |
|  |  | FW3.2 | LW31640 | LW00831 |  |
|  |  | FW11.3 | LW06501 |  |  |
|  |  | OVATE | LW03107 |  |  |
|  |  | SUN | LW34672 |  |  |
|  |  | SlIMA | LW03846 |  |  |
| Cell expansion | Endocycle control | SlCCS52A | LW31221 | LW39104 |  |
|  |  | SlWEE1 | LW20232 |  |  |
|  |  | SlKRP1 | LW19766 |  |  |
|  | Auxin signaling | SlIAA17 | LW16871 |  |  |
|  | ABA biosynthesis | NCED1 | LW01716 | LW05060 | LW29632 |
|  |  | FLACCA | LW05110 |  |  |
|  | Primary metabolism | HXK1 | LW28269 | LW24825 |  |
|  |  | SuSY | LW25002 | LW28024 | LW19118 |
|  |  | LIN5 | LW42636 |  |  |
|  |  | TIV1 | LW31118 | LW42849 |  |
|  |  | mMDH | LW01653 | LW04934 |  |
|  | Regulation of primary metabolism | SPA | LW04513 |  |  |
|  | Ascorbate biosythesis | Gal-LDH | LW42226 |  |  |
|  |  | GME | LW13756 | LW16865 |  |

**Supplementary Table 14 Hub genes list of turquoise module.**

| **Top 10 in network Sheet1 ranked by MCC method.** | | | |
| --- | --- | --- | --- |
| Rank | Name | Score | Annotation |
| 1 | LW19784 | 2954 | LBD10 |
| 2 | LW11286 | 2496 | RPG1 |
| 3 | LW00359 | 1905 | RBOHE |
| 4 | LW09407 | 1104 | CALS5 |
| 5 | LW30895 | 513 | SK32 |
| 6 | LW39354 | 452 | MYB33 |
| 7 | LW42206 | 20 |  |
| 7 | LW23591 | 20 |  |
| 7 | LW01329 | 20 |  |
| 7 | LW42550 | 20 |  |

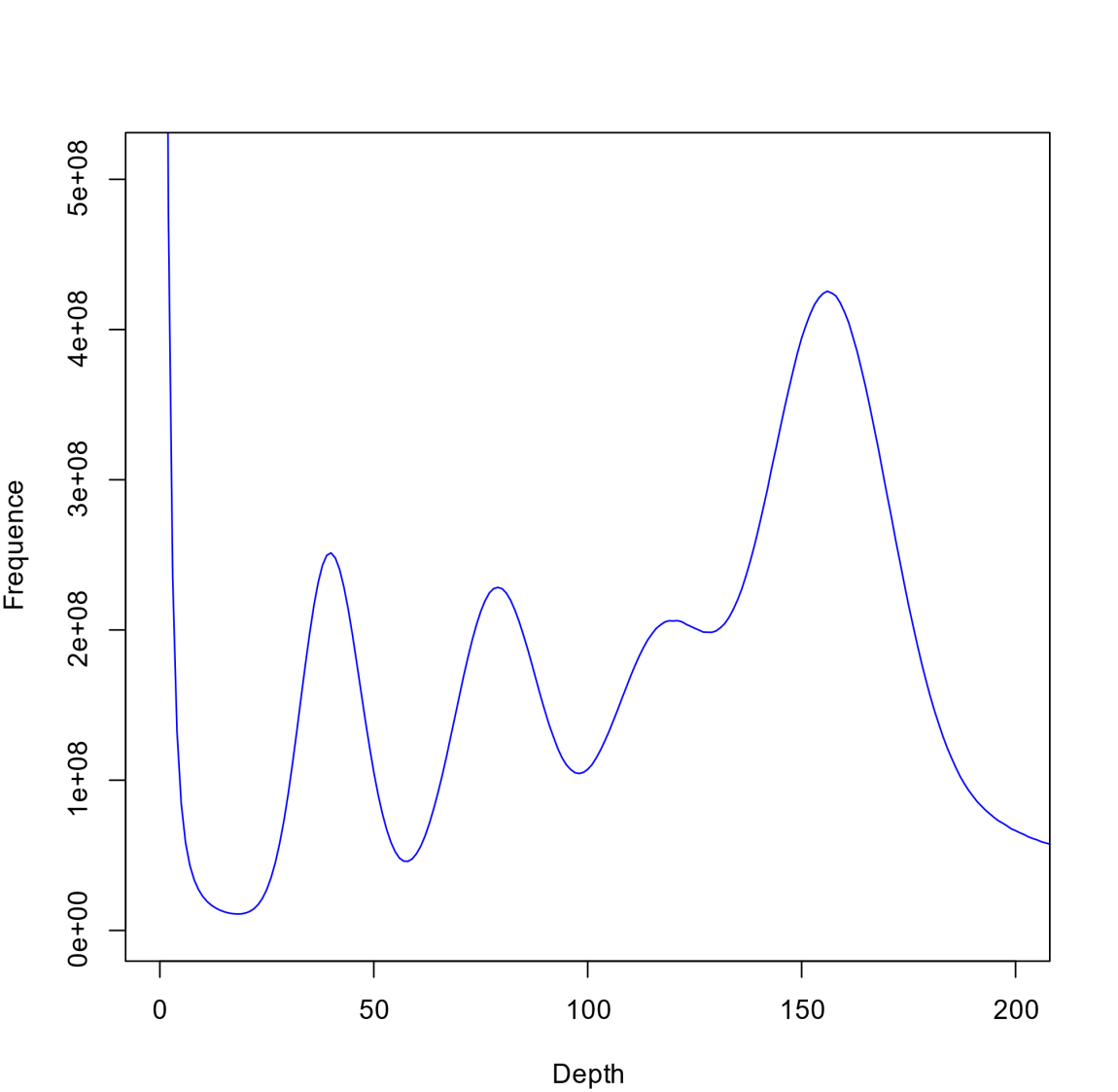

**Supplementary Figure 1 K-mer (17-mer) analysis and estimation of genome size.**

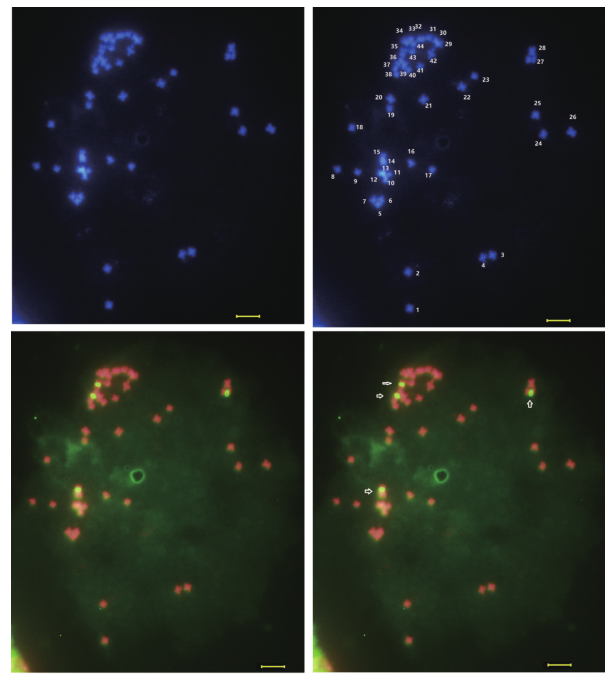

**Supplementary Figure 2 Chromosome karyotypes in ‘Tub’.**

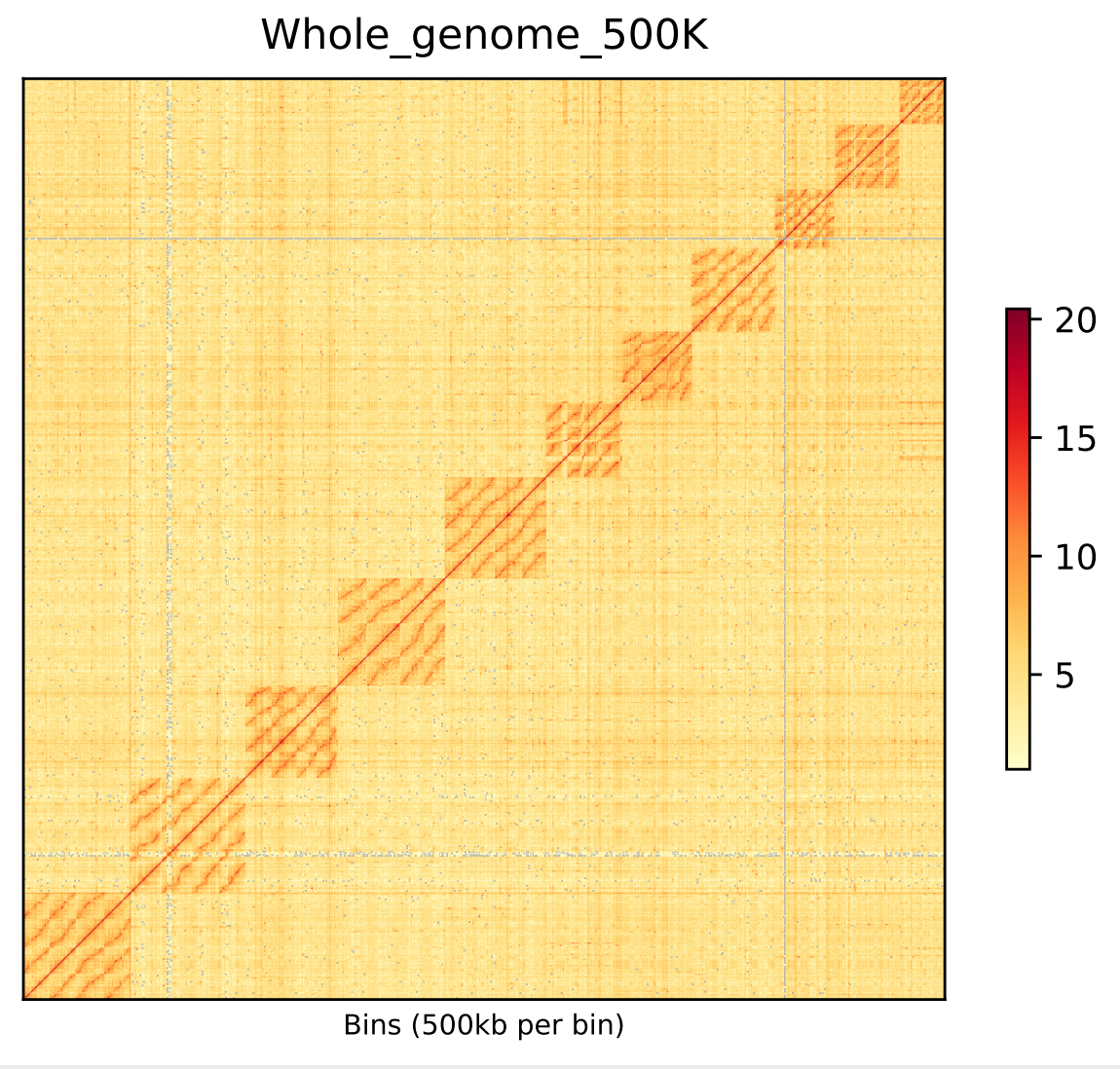

**Supplementary Figure 3 The chromatin interactions of *S. samarangense* revealed 44 assembled chromosomes.**

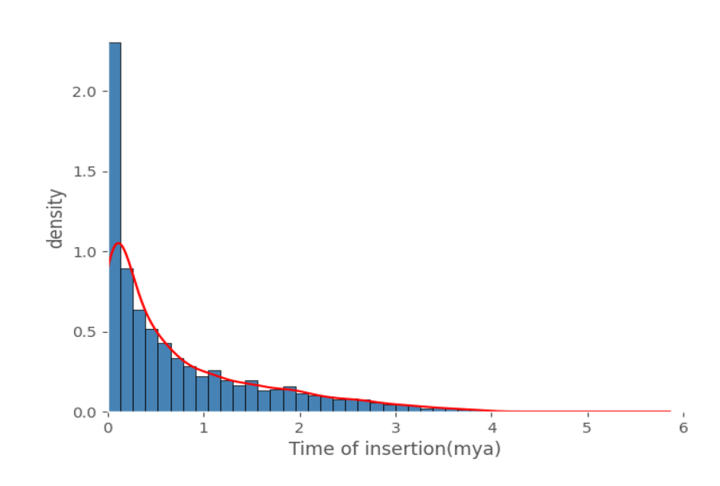

**Supplementary Figure 4 Prediction of the LRT-RT insertion burst time.**

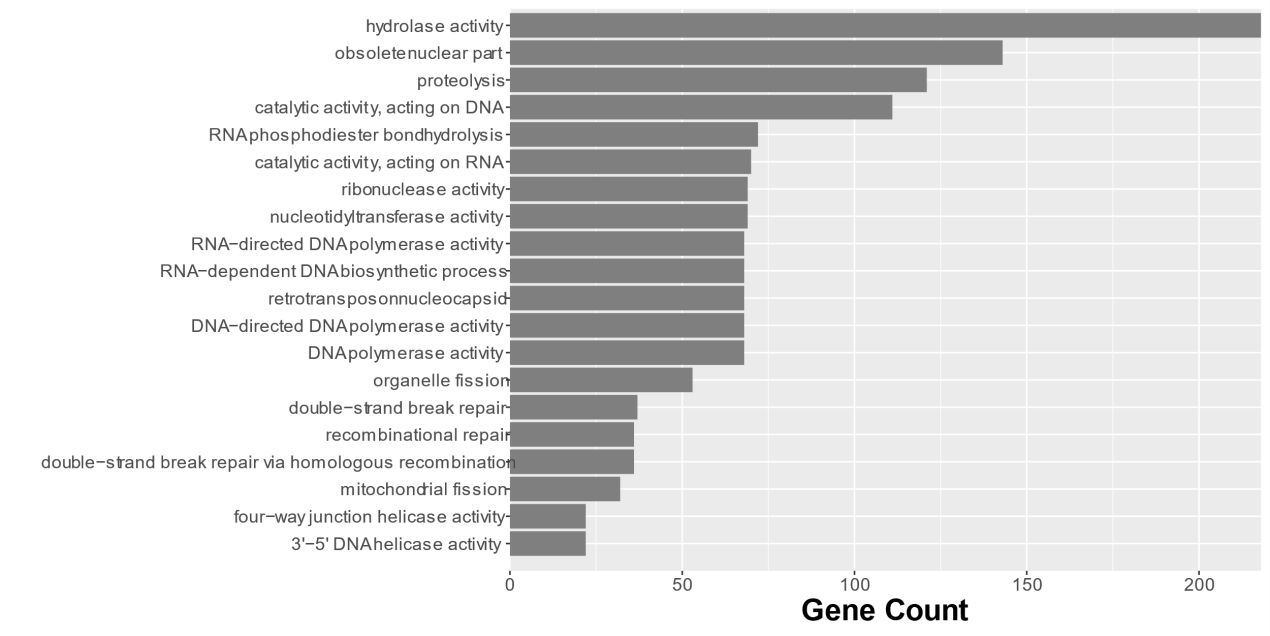

**Supplementary Figure 5 GO enrichment of expansion gene families in *S. samarangense* genome.**

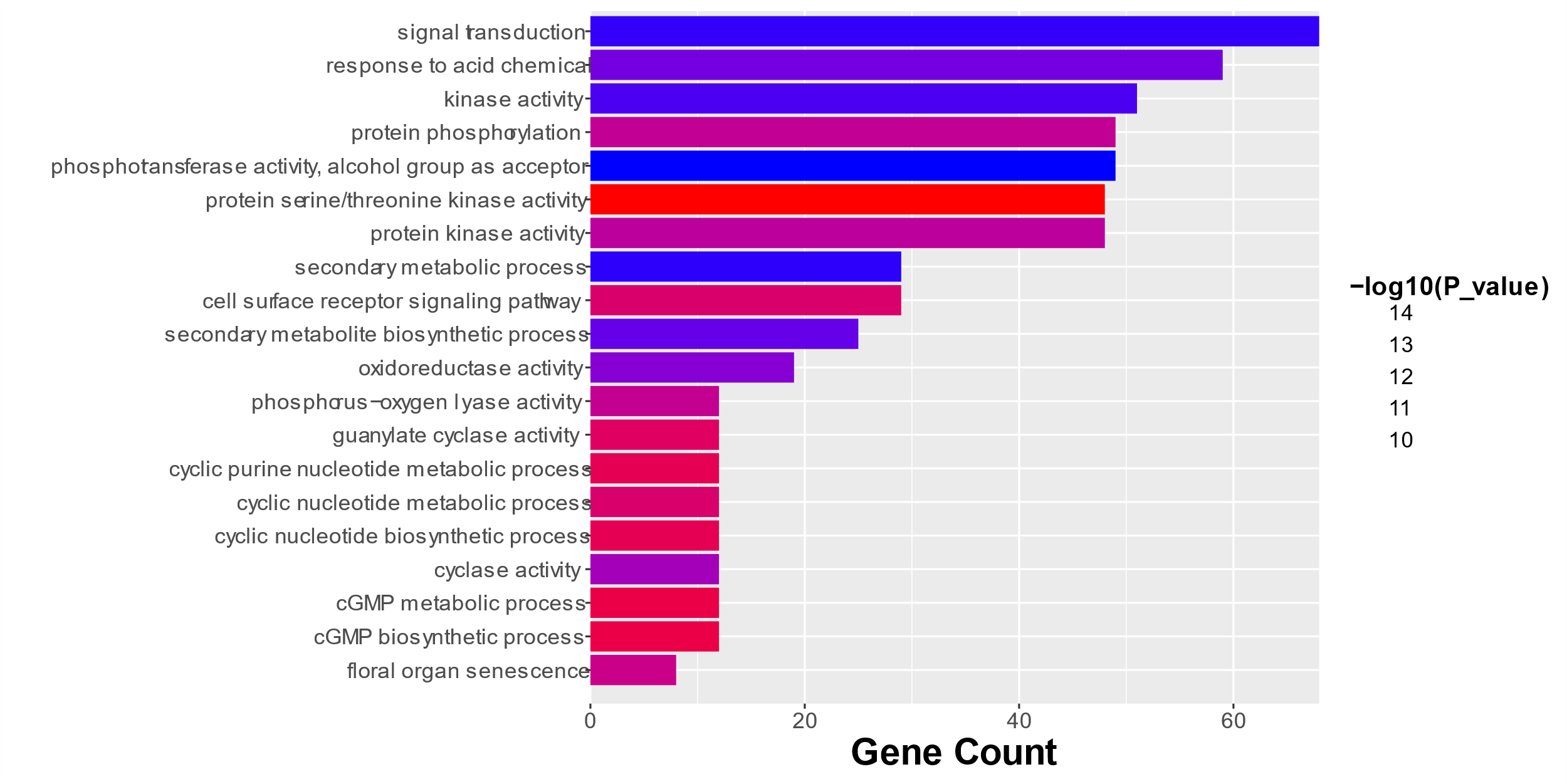

**Supplementary Figure 6 GO enrichment of contraction gene families in *S. samarangense* genome.**

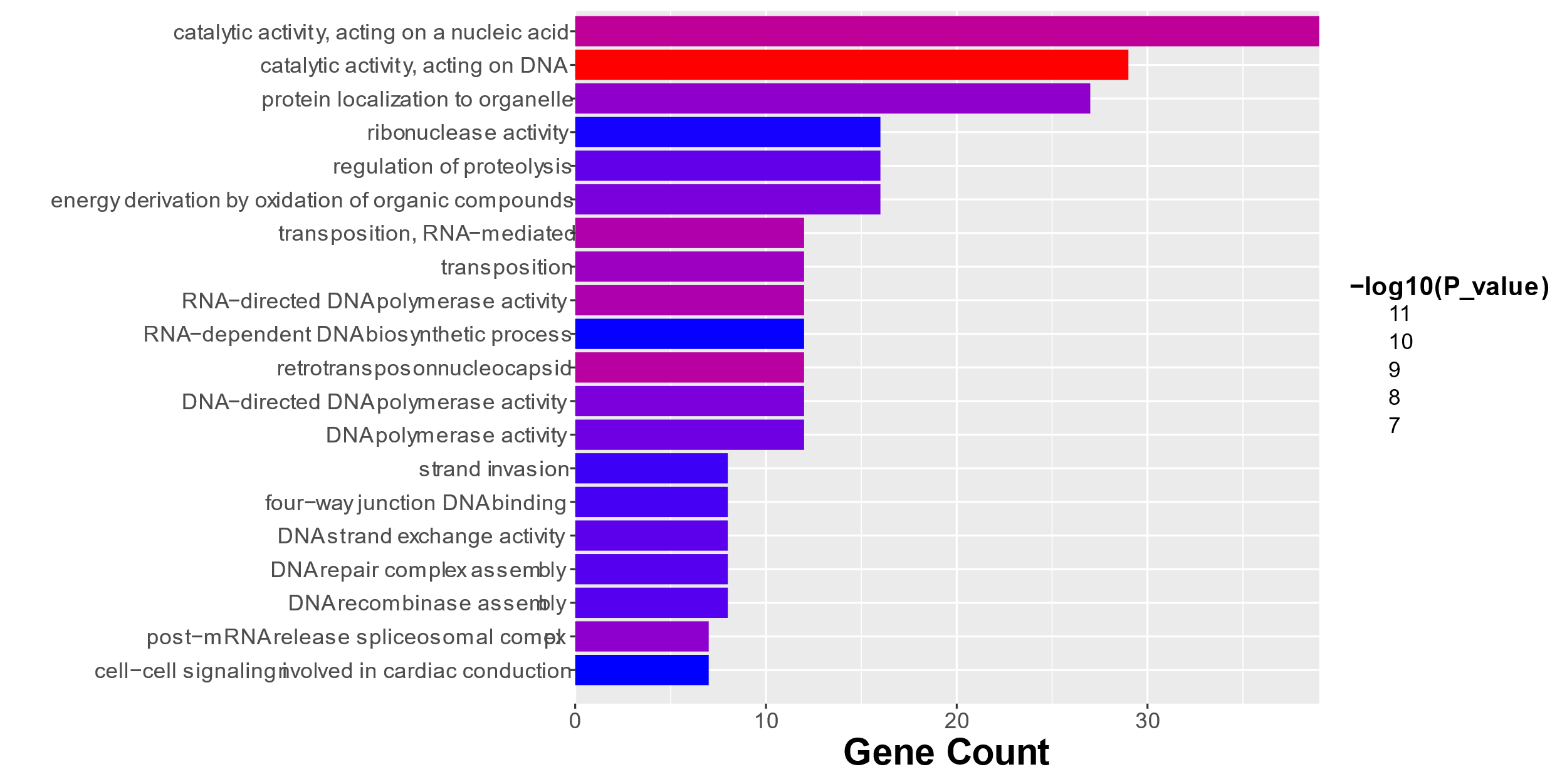

**Supplementary Figure 7 GO enrichment of specific gene families in *S. samarangense* genome.**

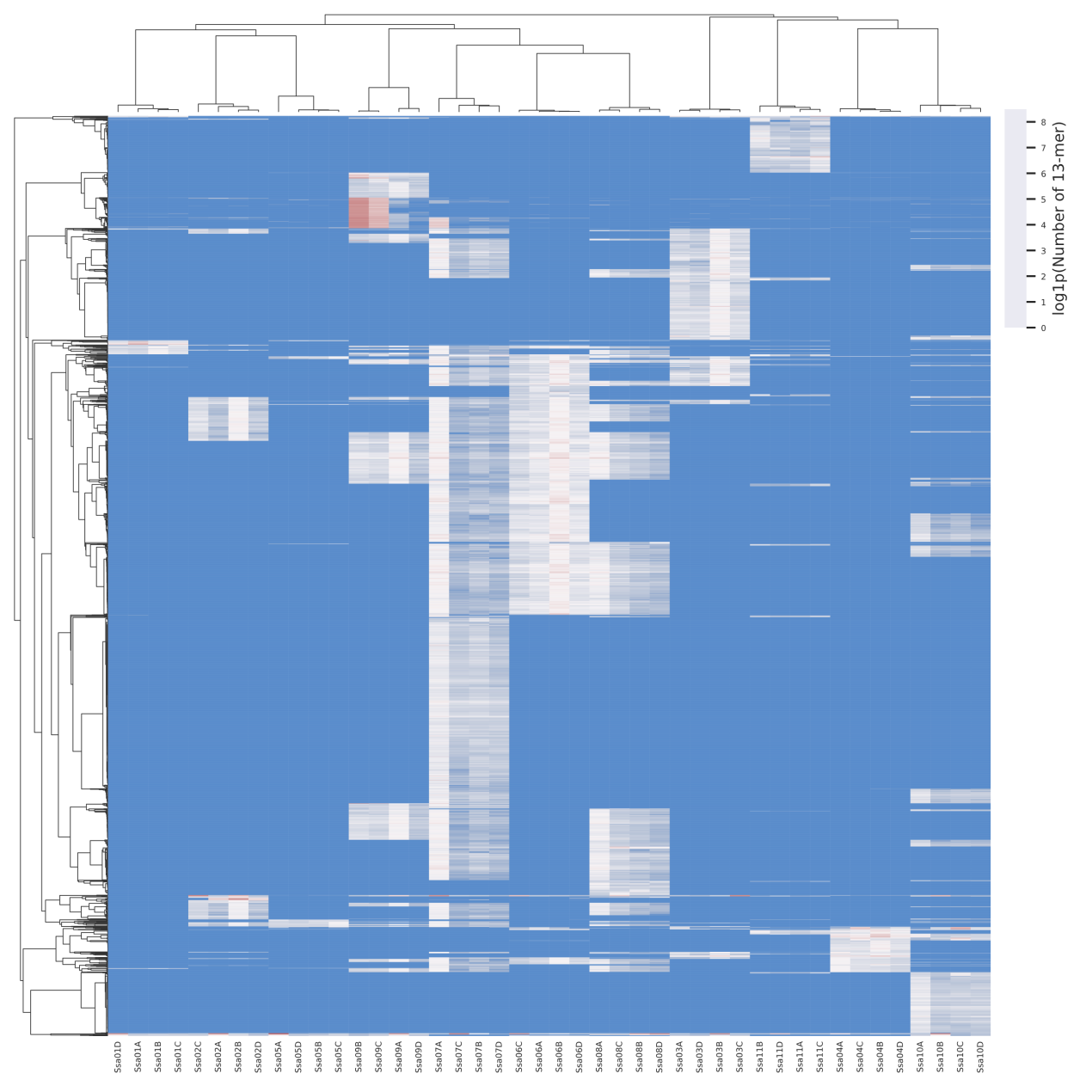

**Supplementary Figure 8 Clustering of 44 chromosomes in the four haplotypes based on the distribution of repetitive elements on chromosomes.**

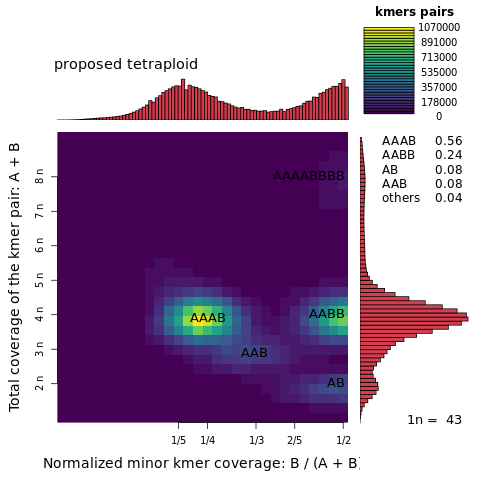

**Supplementary Figure 9 The smudge plot analysis in *S. samarangense.***

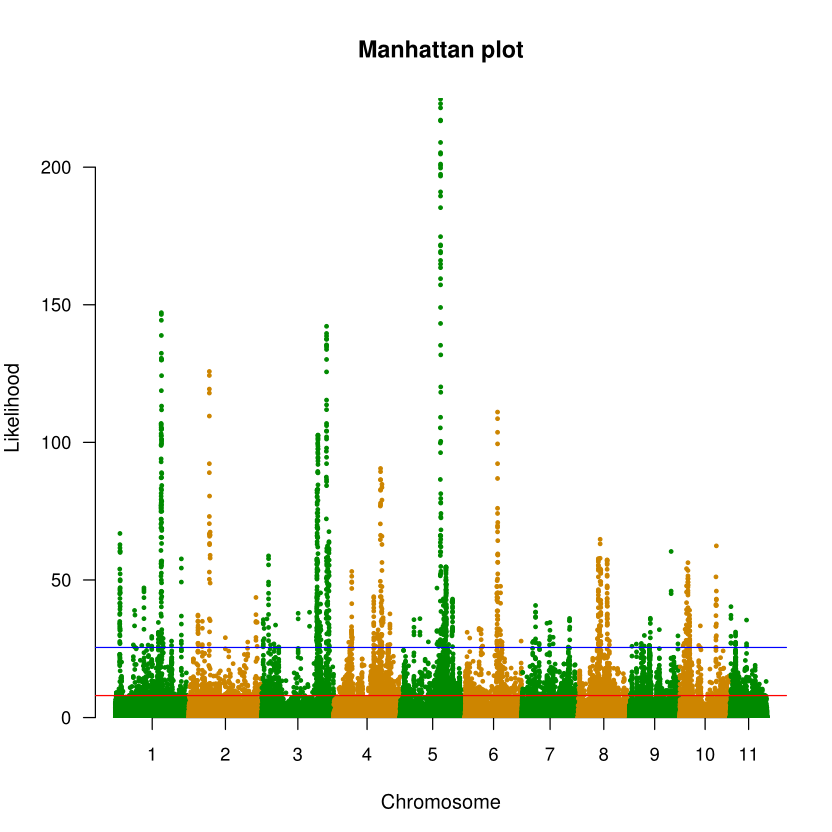

**Supplementary Figure 10 The distribution of selective-sweep signals identified based on SweeD analysis of 27 re-sequenced accessions along 11 chromosomes in group1. The blue line represents the threshold of the top 1%. The red represents the threshold of the top 5%.**

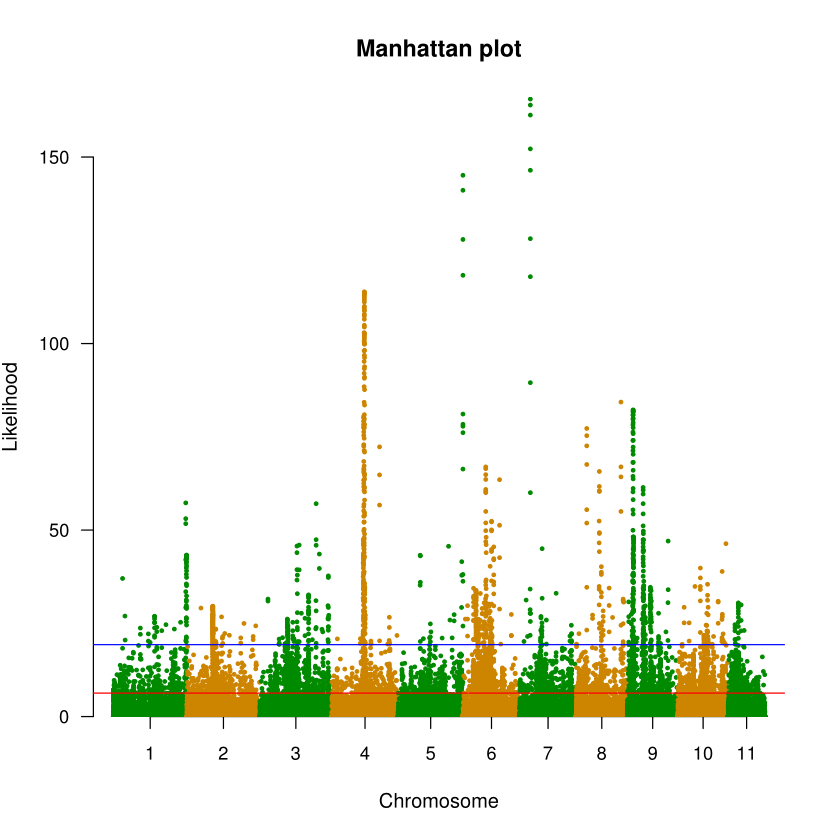

**Supplementary Figure 11 The distribution of selective-sweep signals identified based on SweeD analysis of 27 re-sequenced accessions along 11 chromosomes in group2. The blue line represents the threshold of the top 1%. The red represents the threshold of the top 5%.**

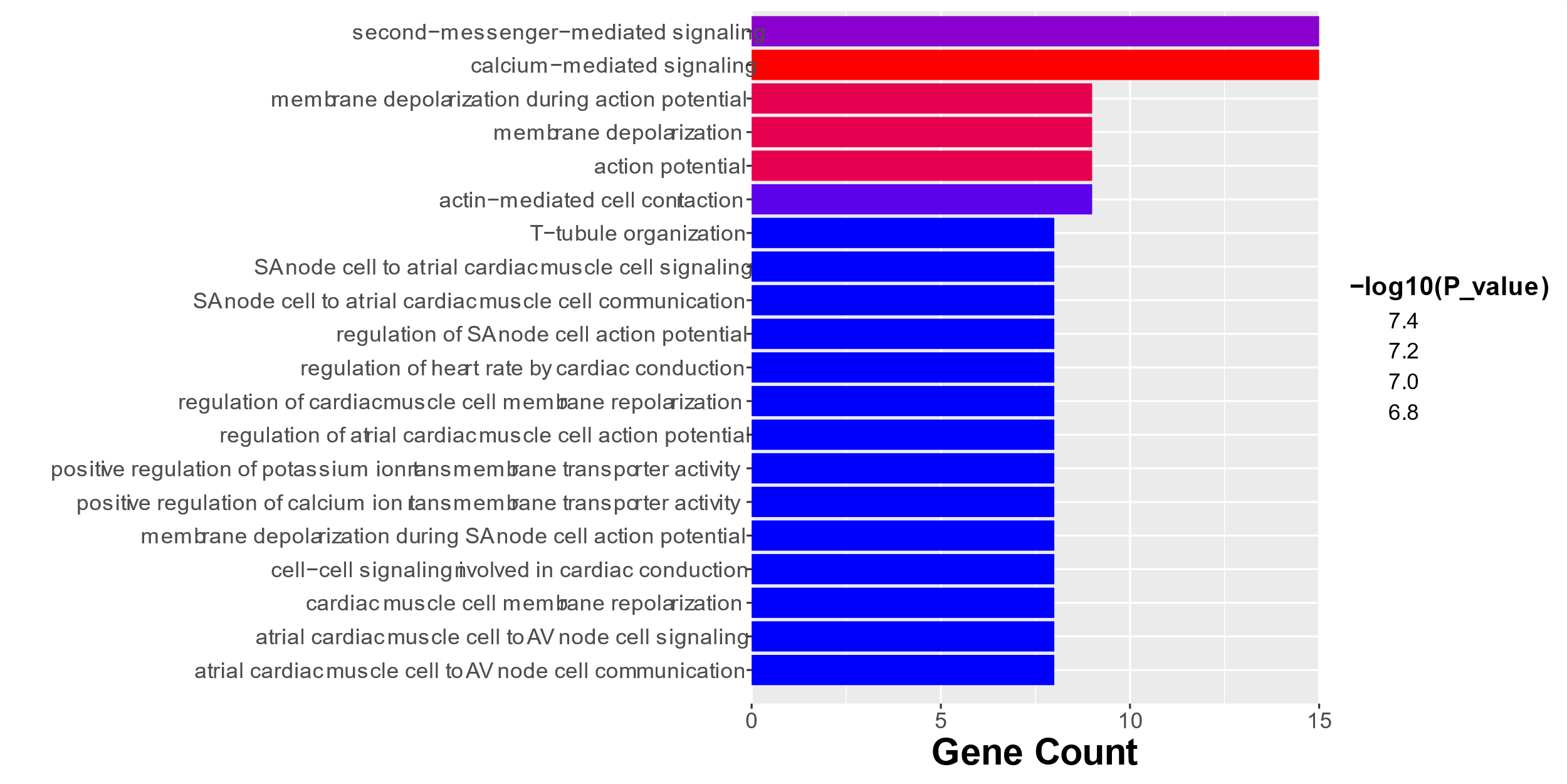

**Supplementary Figure 12 GO enrichment of the swept genes in group1.**

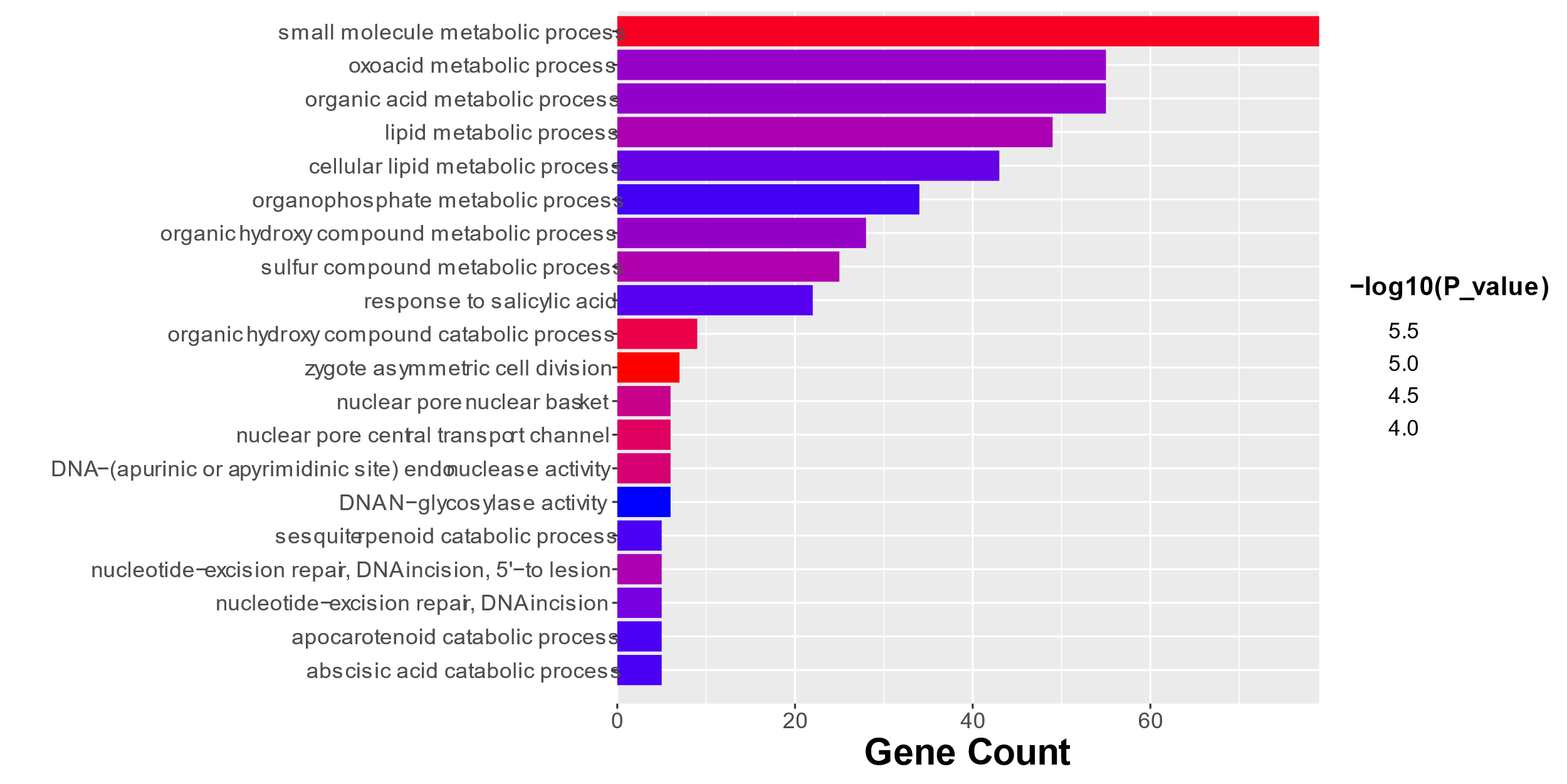

**Supplementary Figure 13 GO enrichment of the swept genes in group2.**

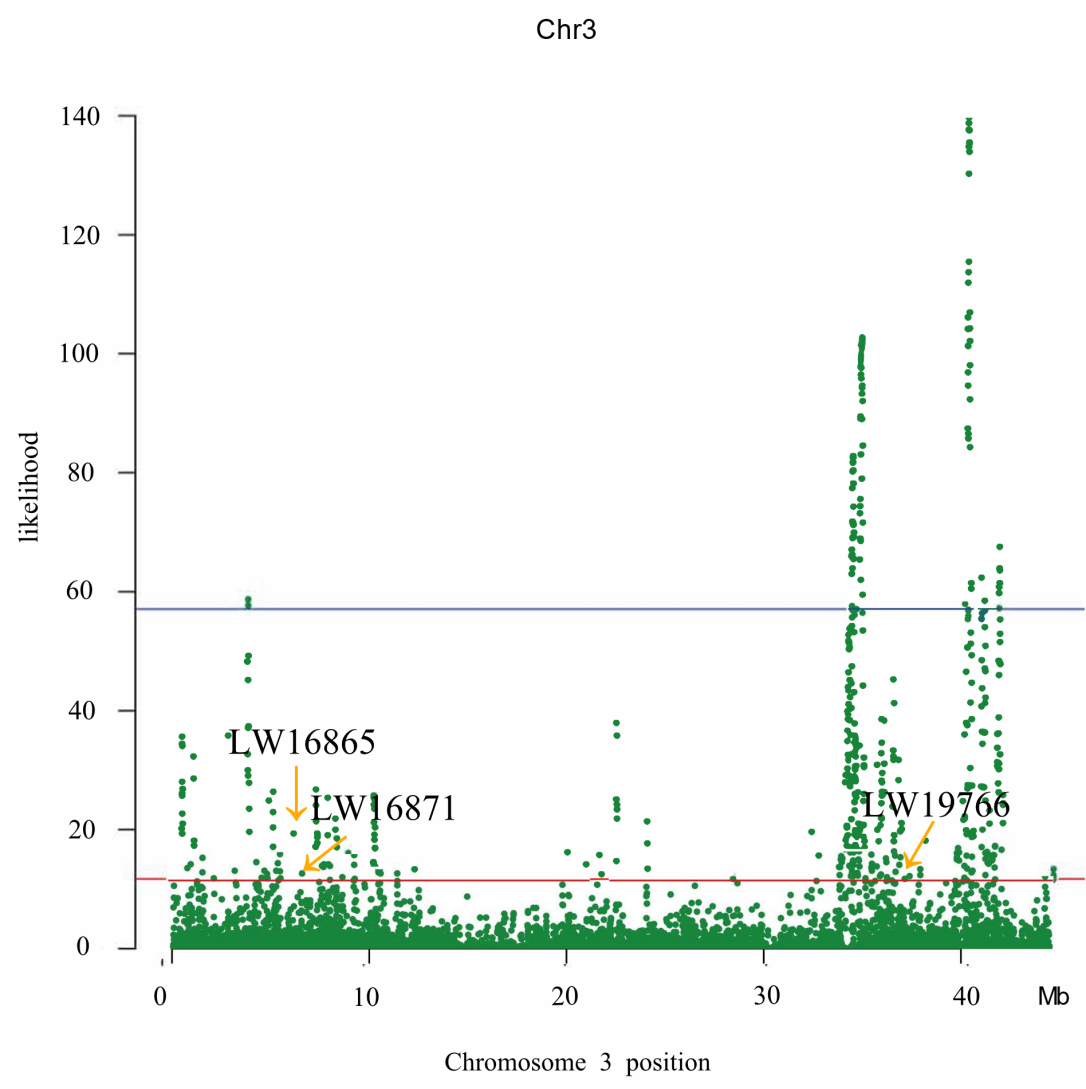

**Supplementary Figure 14 Genes related to fruit growth were identified based on SweeD analysis between landraces and cultivars. Signals of swept region in landraces. The blue line represents the threshold of the top 1%. The red represents the threshold of the top 5%.**

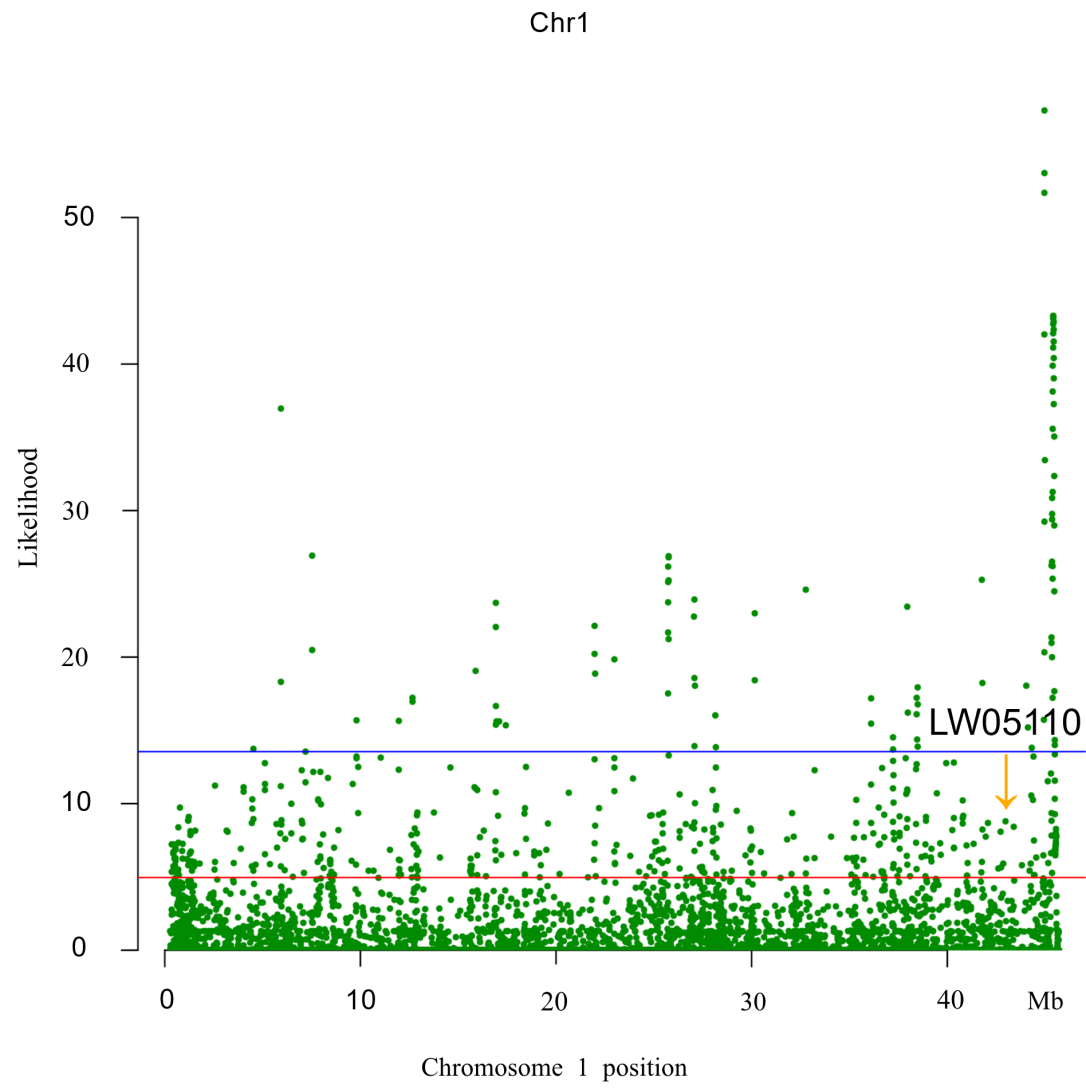

**Supplementary Figure 15 Genes related to fruit growth were identified based on SweeD analysis between landraces and cultivars. Signals of swept region in cultivars. The blue line represents the threshold of the top 1%. The red represents the threshold of the top 5%.**

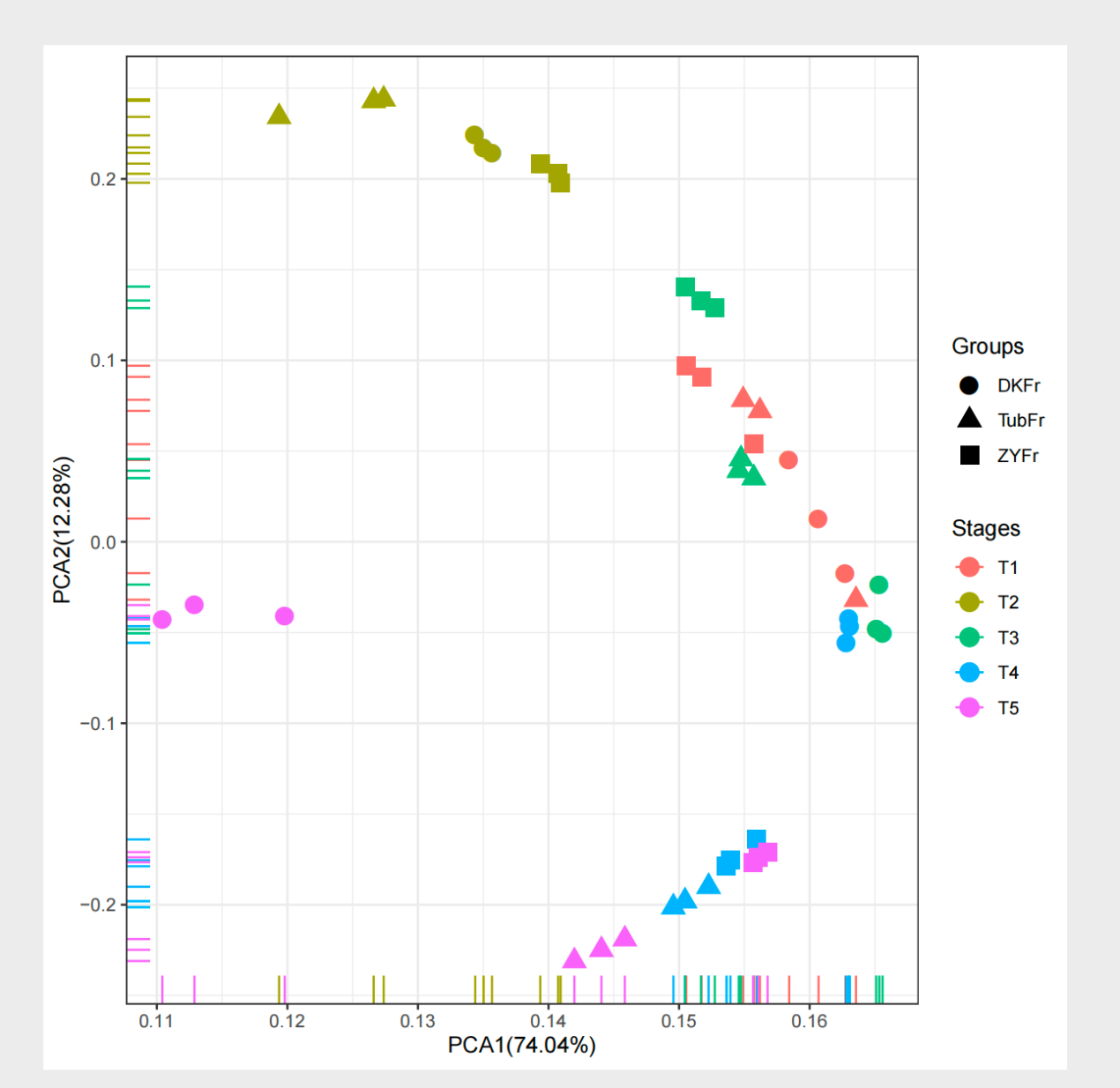

**Supplementary Figure 16 Principal component analysis (PCA) of the different fruit development transcriptome data.**

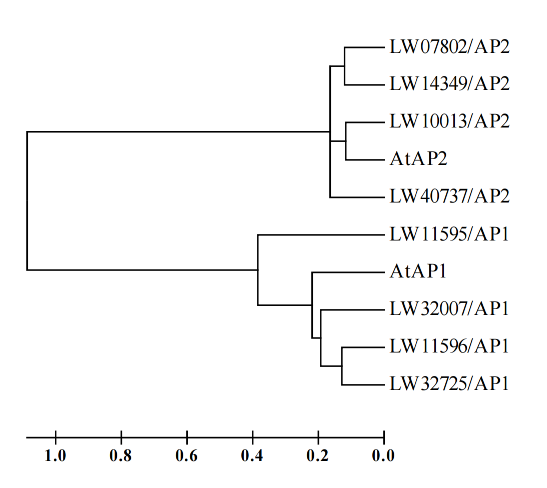

**Supplementary Figure 17 Phylogenetic tree of ten genes of AP1 and AP2 from wax apple and *Arabidopsis thaliana*. The tree was constructed with MEGA4 software.**

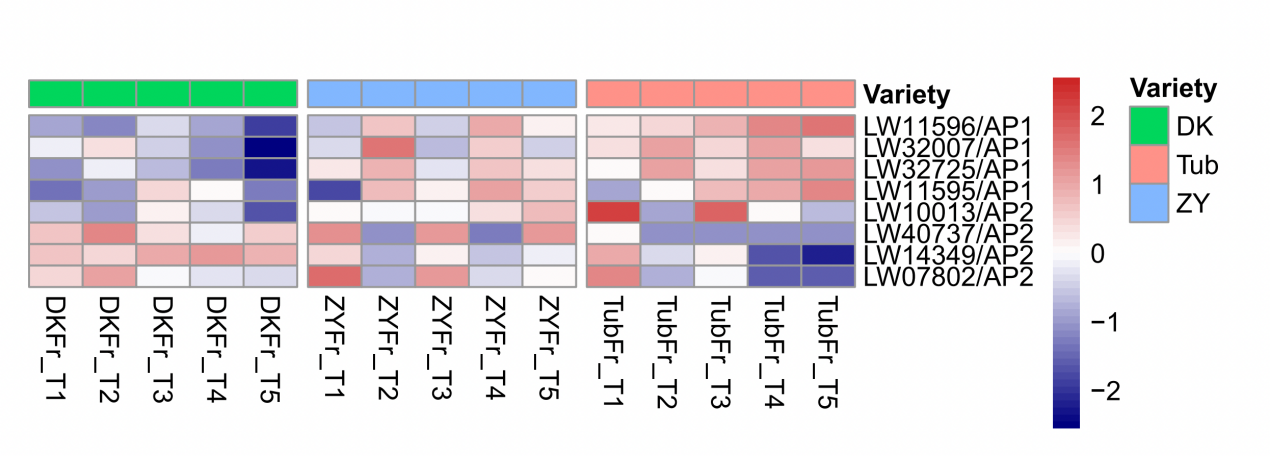

**Supplementary Figure 18 Expression (FPKM) of eight AP1 and AP2 genes in ‘DK’, ‘ZY’, and ‘Tub’ from different fruit development stages. The expression from low to high is indicated by the scale ranging from blue to red.**

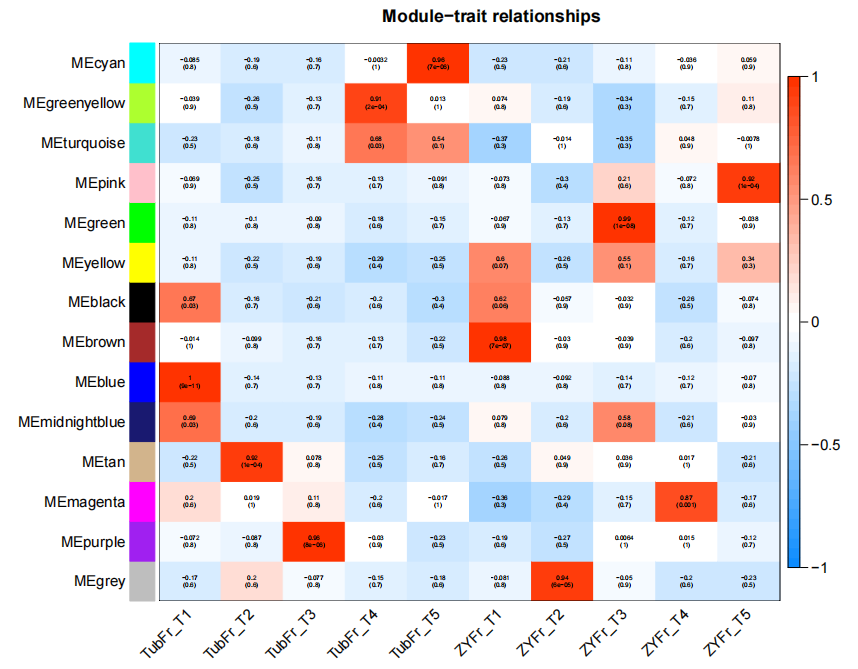

**Supplementary Figure 19 Pearson correlation coefficient between module and sample feature based on fruit development analysis. Each row represents a different module and numbers within the modules represent correlation coefficients and P values. The color scale at the right indicates correlation coefficients.**

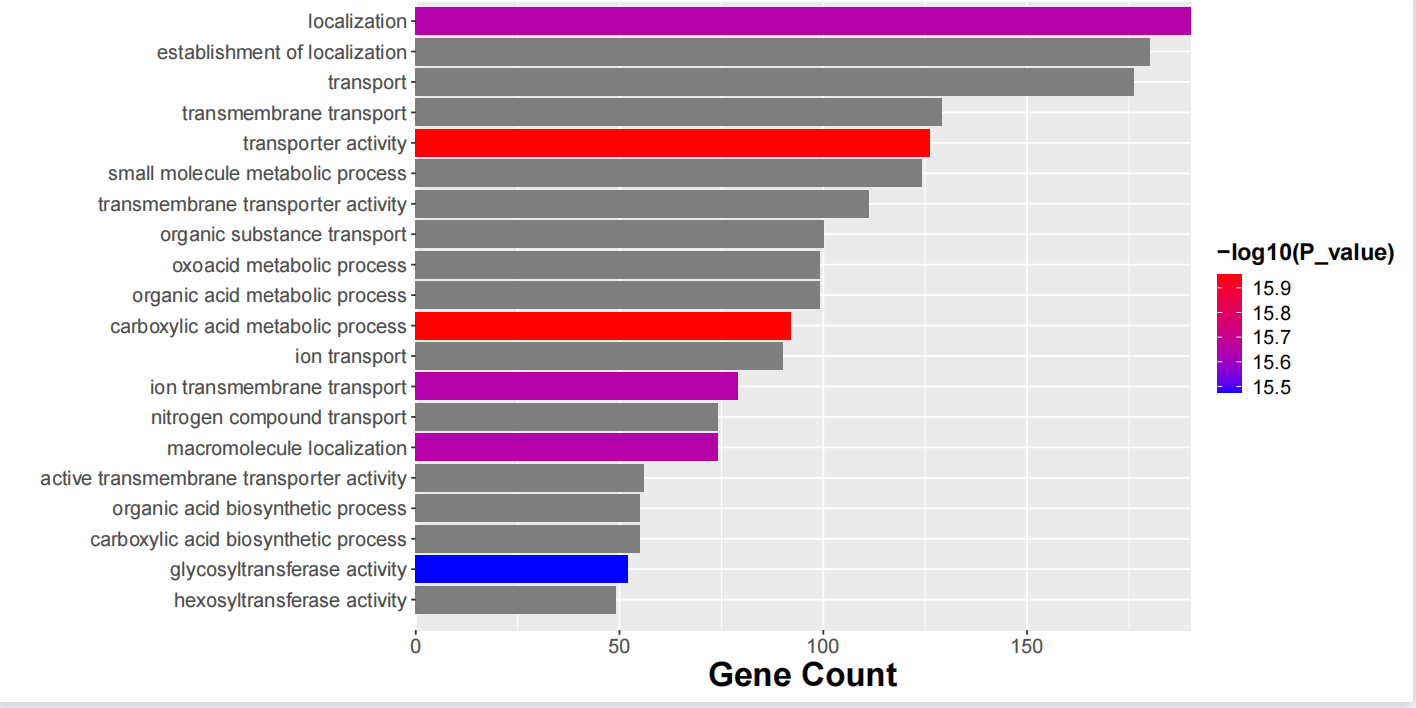

**Supplementary Figure 20 GO enrichment of the genes in pink module.**

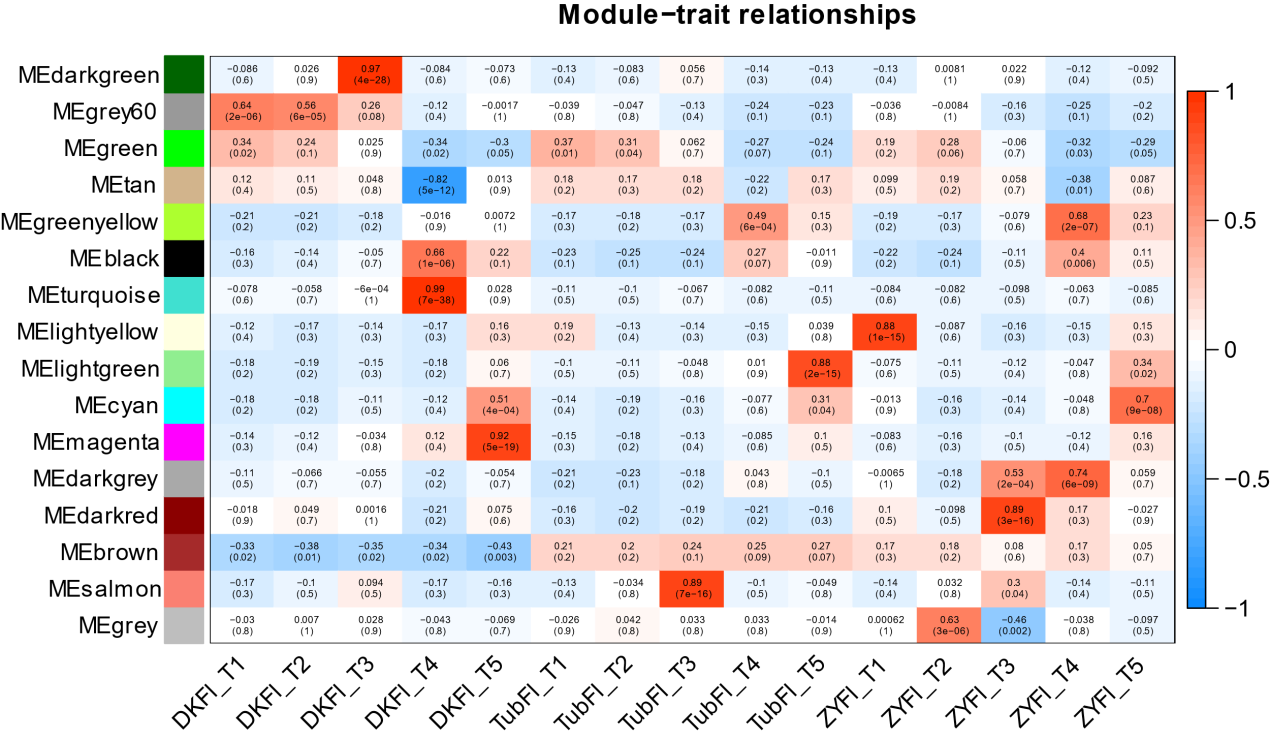

**Supplementary Figure 21 Pearson correlation coefficient between module and sample feature based on flower development analysis. Each row represents a different module and numbers within the modules represent correlation coefficients and P values. The color scale at the right indicates correlation coefficients.**

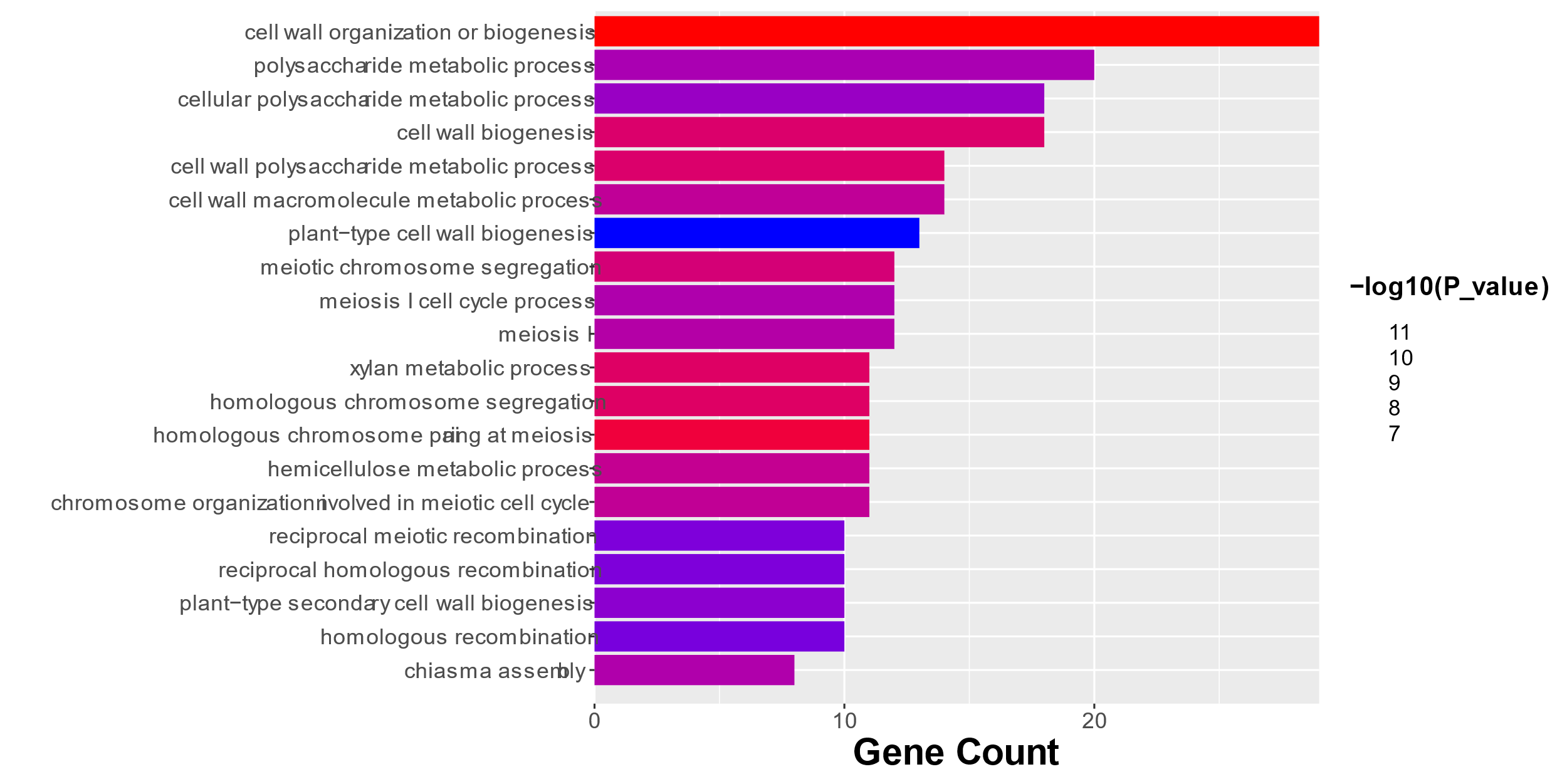

**Supplementary Figure 22 GO enrichment of darkgreen module.**

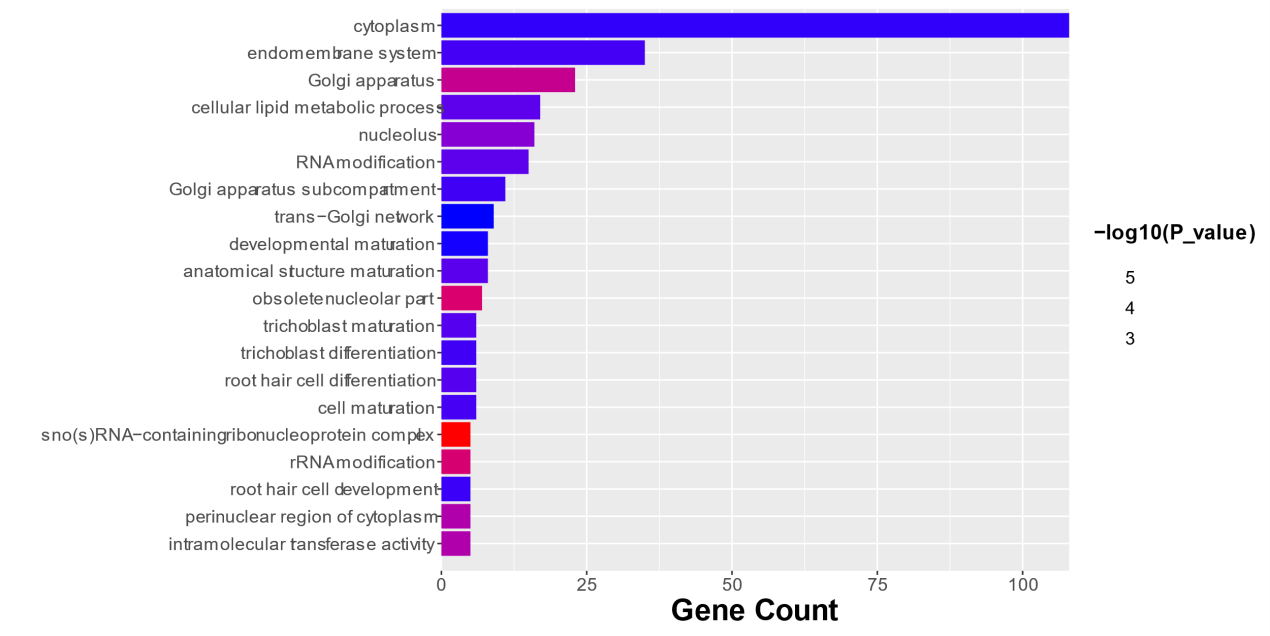

**Supplementary Figure 23 GO enrichment of tan module.**

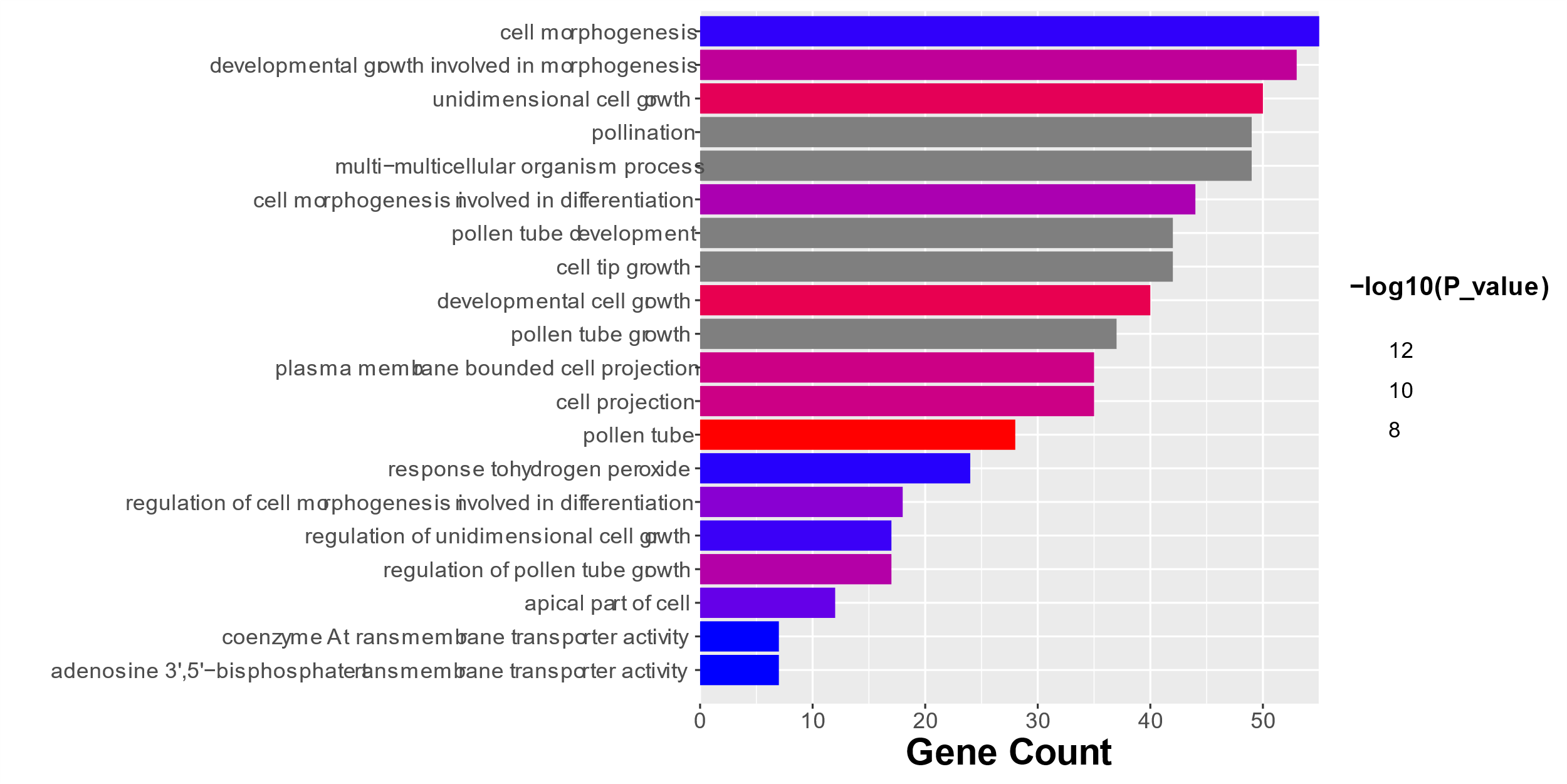

**Supplementary Figure 24 GO enrichment of turquoise module.**

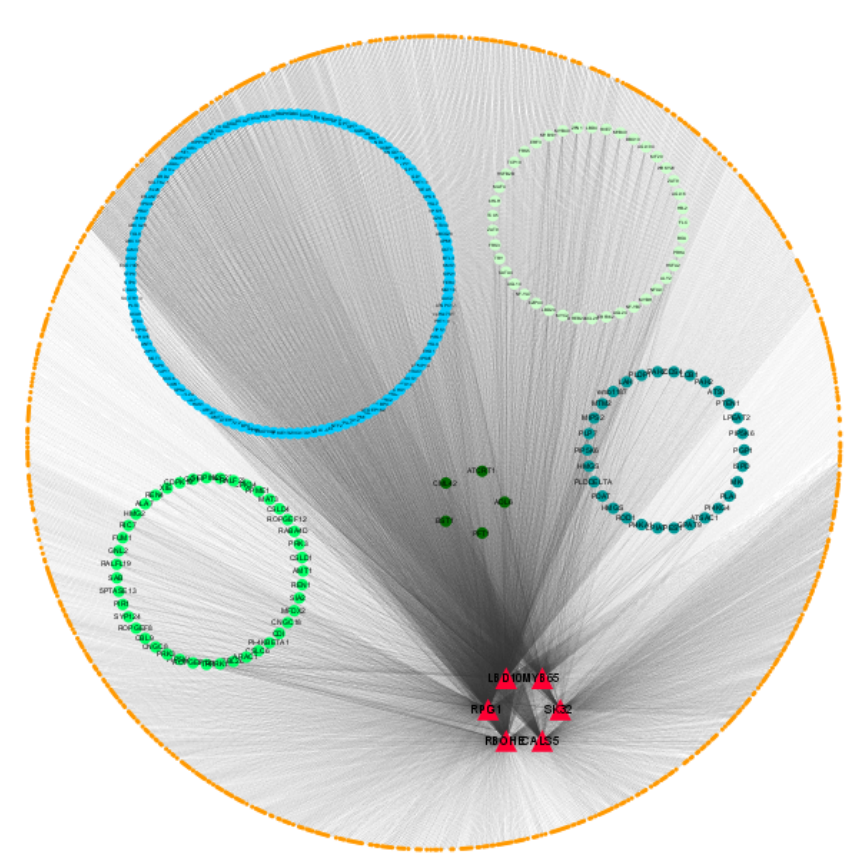

**Supplementary Figure 25 The correlation networks in the turquoise module. Candidate hub genes are shown in red.**

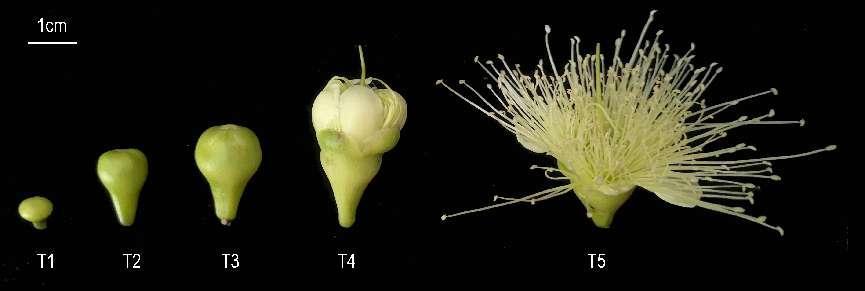

**Supplementary Figure 26 The flowers from three varieties including ZY, Tub, and DK were sampled from T1 to T2.**
